# The ultrasmall ocean microbiome: a reservoir of microbial diversity and nitrogen fixation

**DOI:** 10.64898/2026.06.16.732551

**Authors:** C.A. Höllerer, S. Miravet-Verde, L.J. Ustick, P. Bork, S.G. Acinas, E. Pelletier, S. Sunagawa, C. Bowler

## Abstract

Marine microbiology has long considered ultrasmall (<0.2 µm) size fractions as virus-dominated, yet emerging evidence suggests potential prokaryotic activity with implications for global biogeochemical cycles. Here, we present the first comprehensive genome-resolved survey of this fraction, built on 4,058 metagenome-assembled genomes (MAGs) representing 1,152 species across Bacteria (90.7%) and Archaea (9.3%), dominated by Pseudomonadota, Bacteroidota, and Nanoarchaeota. Over 490 non-redundant MAGs occurred exclusively in the ultrasmall fraction, with 66.8% representing novel taxa, particularly abundant in Arctic surface waters. Beyond known ultrasmall organisms (e.g., DPANN archaea and Patescibacteria), which show expected genomic and metabolic reduction, functional annotation revealed diverse metabolic potential across nitrogen, sulfur, and carbon cycling. Focusing on nitrogen fixation, we identified 13 non-cyanobacterial diazotroph species with complete nitrogenase operons spanning four bacterial classes. These resolved into two ecotypes: cosmopolitan low-abundance taxa, and Arctic ultrasmall-restricted strains reaching up to 16.7% relative abundance. Two lineages (*50-400-T64* and *Novosphingobium)* showed ultrasmall exclusive bipolar distributions across Arctic and Southern Oceans. Beyond nitrogen fixation, these diazotrophs encoded pathways for denitrification, DNRA, sulfur oxidation, and carbon fixation. Our findings reveal an overlooked reservoir of microbial diversity including diazotrophs in the ultrasmall ocean microbiome, with significant implications for polar nitrogen budgets and global biogeochemical models.

## Introduction

Marine microbial communities are among the most diverse biological systems on Earth that collectively drive global biogeochemical cycles and primary productivity^1^. As the vast majority of microorganisms remain uncultivated, culture-independent approaches based on metagenomes and, in particular, metagenome-assembled genomes (MAGs) offer significant new insights into their diversity, distribution and functional potential^2–4^. Marine samples are often separated by size fractions to reflect the different ecological niches: free-living organisms, particle-attached or aggregates, and the ultrasmall size fraction^5,6^. While the ultrasmall size fraction was long believed to be restricted to viruses, two major prokaryotic radiations, the DPANN archaea (Diapherotrites, Parvarchaeota, Aenigmarchaeota, Nanoarchaeota, and Nanohaloarchaeota) and the Patescibacteria (also known as Candidate Phyla Radiation, CPR), are both known to include lineages with ultrasmall cell sizes (100–400 nm)^7,8^, and are characterized by highly reduced genomes and symbiotic or host-dependent lifestyles^7–13^. Despite their genomic reduction, some ultrasmall microorganisms participate in key metabolic processes including carbon fixation, carbon and sulfur cycling^14–17^.

Nitrogen is an essential macronutrient in marine ecosystems^1,18^, yet approximately 95% exists as inert dinitrogen gas (N_2_), which is inaccessible to most organisms^19^. Biological nitrogen fixation (BNF), the conversion of atmospheric N_2_ into bioavailable forms by specialized microorganisms known as diazotrophs, is a critical source of new nitrogen, particularly in oligotrophic ocean regions^20–22^. Historically, microscopy-based observations established filamentous cyanobacteria such as *Trichodesmium* as the dominant marine diazotroph in warm, oligotrophic waters^23^. Consistent with this, direct rate measurements and inverse biogeochemical models show that BNF is highest in low latitude regions^24,25^ and decreases towards higher latitudes^26,27^.

Diazotrophs possess the nitrogenase enzyme complex, whose iron protein (*nifH*) serves as a widely used molecular marker^28^. Molecular surveys, targeting *nifH*, drastically increased the known diversity of diazotrophs^3,28^, though *nifH* phylogeny alone provides limited taxonomic resolution and does not confirm functional diazotrophy potential without the minimal nitrogenase cluster set (*nifH/D/K*)^3^. Genome-resolved metagenomics have addressed these limitations, enabling more robust taxonomic assignment and reconstruction of metabolic potential^29,3,30^. These approaches revealed diverse diazotrophs, including unicellular cyanobacteria and non-cyanobacterial diazotrophs (NCDs) from bacterial and archaeal lineages, which dominate diazotroph communities in some ocean regions^29,31^ and display lifestyles ranging from particle-associated to free-living and potentially symbiotic^29,31^. Metagenomic surveys detecting diazotroph genes suggest potential nitrogen fixing activity in additional polar regions^32,33^ and deep-sea environments^34,35^. Surprisingly, recent studies also report evidence of nitrogen fixation potential in the ultrasmall size fraction, with particularly high occurrences in the Arctic Ocean^32,36^.

The detection of *nifH* genes in the ultrasmall size fraction suggests a potentially undiscovered niche for marine diazotrophs. However, the validation of functional diazotrophy, along with the taxonomic identity of these *nifH* hosts and their metabolic potential, remain unknown without genome-resolved characterization.

In this study, we explored over 4,000 metagenome-assembled genomes (MAGs) recovered from ultrasmall size fraction (<0.2 um) metagenomes retrieved from the Ocean Microbiomics Database (OMDB, omdb.microbiomics.io), which represents over 12,000 globally distributed samples, of which 315 samples correspond to the size fraction <0.2 um. The ultrasmall fraction samples were processed to enrich prokaryotic cells while removing viral sequences. We provide a comprehensive analysis of ultrasmall prokaryotic communities, including their taxonomic diversity, distribution and metabolic potential, and specifically focus on putative diazotrophs.

## Results and Discussion

### Diversity and distribution of MAGs from the ultrasmall size fraction

#### General overview of over 4,000 MAGs from the ultrasmall size fraction

A total of 4,058 metagenome-assembled genomes (MAGs) originating from the viral size fraction (0–0.2 μm) were retrieved from the OMDB (omdb.microbiomics.io). These MAGs were compiled from nine independent studies (Fig. 1a, Supplementary Table S1)^37–45^, covering all major ocean basins, with sampling spanning both epipelagic and mesopelagic depths. Average nucleotide identity (ANI) clustering resulted in a set of 1,152 non-redundant species (95% ANI) and 1,824 strain-level clusters (99% ANI). Genome completeness across the dataset was high, with an average of 86% and low contamination values averaging 1.24 % (Fig. 1d).

**Figure 1.**
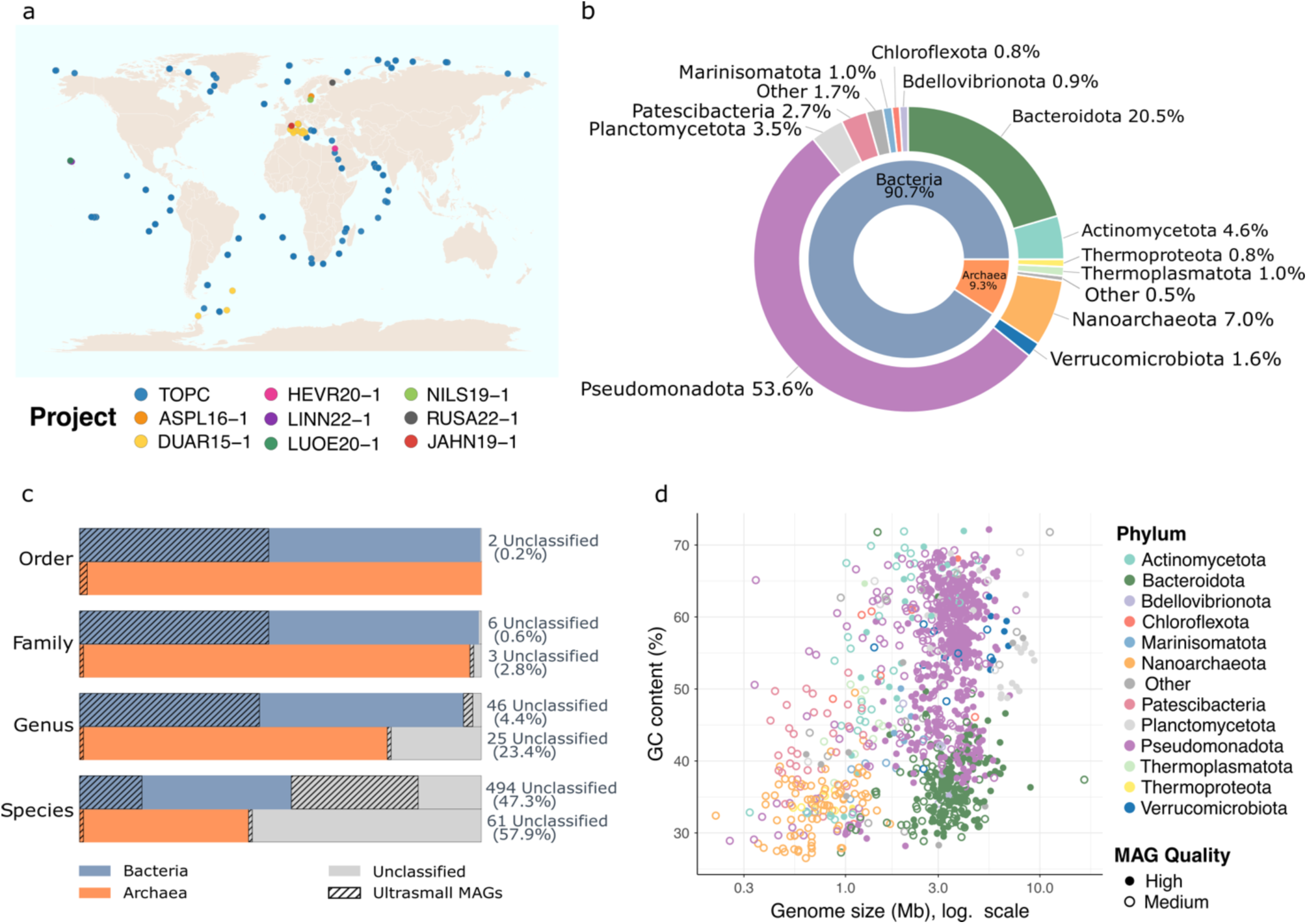
Overview of MAGs from ultrasmall size fractions. **a.** Global distribution of sampling sites from projects contributing MAGs to the ultrasmall size fraction dataset. **b.** Taxonomic composition based on GTDB annotations. The inner ring denotes domain-level assignments, while the outer ring shows corresponding phylum-level classifications. **c.** Taxonomic resolution across ranks for Bacteria and Archaea. Bars indicate the proportion of clusters with resolved taxonomy per rank. Exact percentages and number of MAGs are shown to the right of the bars. Hashed lines indicate MAGs found exclusively in the ultrasmall size fraction. **d.** Genomic characteristics of representative MAGs at 95% ANI. Each point corresponds to a MAG, plotted by genome size (x-axis) and GC content (y-axis). Points are colored by phylum and styled according to MAG quality (High Quality: ≥90% completeness and <5% contamination, Medium Quality: ≥50% completeness values and <10% contamination).

GTDB taxonomic classification revealed that 90.7% of dereplicated MAGs were bacterial (n=1,045) and 9.3% were archaeal (n=107). At the phylum level, Pseudomonadota was the most abundant (n=617), followed by Bacteroidota (n=236), Nanoarchaeota (n=81), Planctomycetota (n=40) and Patescibacteria (n=31) (Fig. 1b). The most represented classes included the bacterial Gammaproteobacteria (n=313), Alphaproteobacteria (n=304), Bacteroidia (n=224), Actinomycetes (n=29) and archaeal Nanoarchaeia (n=81), Poseidoniia (n=11) and Nitrososphaeria (n=9). Notably, the dataset includes MAGs belonging to the Patescibacteria (former Candidate Phyla Radiation; CPR) and the DPANN superphylum (in this dataset: Aenigmatarchaeota, Iainarchaeota, Nanoarchaeota, Undinarchaeota), known to be associated with ultrasmall cell sizes. Taxonomic novelty was high, with less than 42 % of archaeal and 52 % of bacterial MAGs having an assignment at species level (Fig. 1c).

Two bacterial species could not be assigned beyond the class Paceibacteria (Phylum Patescibacteria), which is a known ultrasmall bacterial lineage. After inspection for likely artifacts (Supplementary Text) MAG genome sizes in our dataset ranged from 0.21 to 11.2 Mb Mb, with a mean of 3.05 ± 1.5 Mb (Fig. 1d) and were strongly structured by phylogeny. MAGs affiliated with the Patescibacteria and DPANN superphyla consistently presented the smallest genome sizes, typically between 0.3 and 1 Mb. These small genomes are a common trait of ultrasmall lineages, and have been linked to extensive metabolic reduction, limited biosynthetic capabilities, and are consistent with symbiotic or host-dependant lifestyles reported in previous studies^11,46^. In contrast, phyla such as Pseudomonadota and Bacteroidota displayed substantially larger genomes, averaging 3.3 ± 1.1 Mb and 3.6 ± 1.4 Mb, respectively, consistent with genome sizes reported for these phyla in larger size fractions^47,48^.

#### Origin and distribution of the ultrasmall size fraction MAGs

The MAGs were recovered from a wide range of ocean regions, with the highest numbers from the Arctic Ocean (42% of MAGs). Archaeal and Patescibacteria MAGs were more frequently assembled from samples collected in the Indian and Pacific Oceans (Supplementary Fig. S1). The NMDS ordination of samples based on MAG composition (Bray-Curtis dissimilarities) revealed a clear structuring of communities by ocean basin and size fraction (Fig. 2a, stress = 0.131). PERMANOVA analysis confirmed ocean basin as the primary driver (R² = 0.087, F = 13.62, p < 0.0001) followed by size fraction (R² = 0.051, F = 21.45, p < 0.0001) (Supplementary Table S2a).

**Figure 2.**
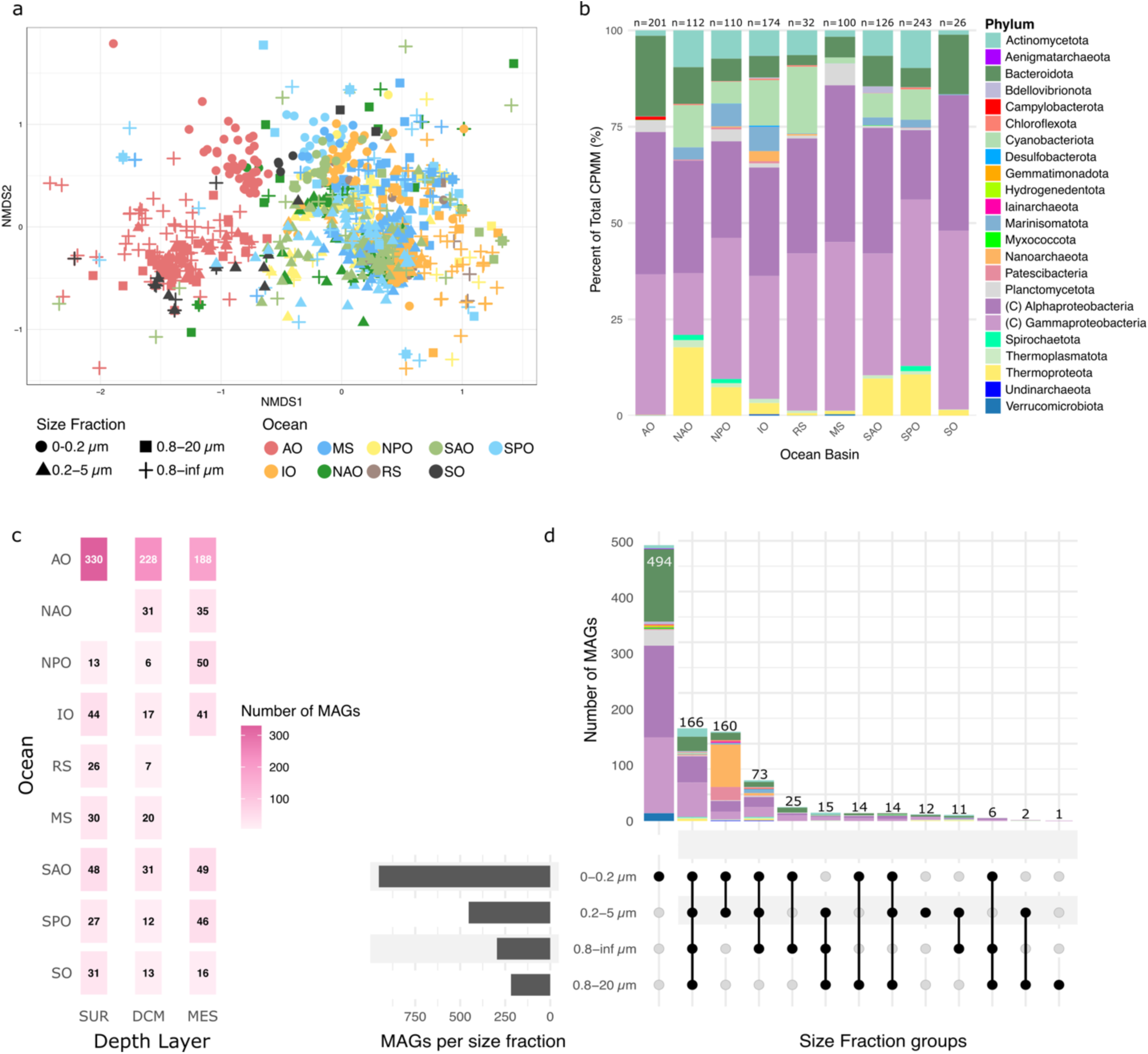
Composition and biogeography of microbial MAGs from ultrasmall size fractions. **a.** Non-metric multidimensional scaling (NMDS) based on Bray-Curtis dissimilarity of MAG abundance profiles. Each point represents a sample, shaped by size fraction and colored by ocean basin. **b.** Stacked bar plots showing the relative abundance of the 1,152 MAGs from the ultrasmall size fraction (phylum level, except for Pseudomonadota, where the Class (C) is shown) in each ocean basin. n indicates the number of samples per ocean. **c.** Heatmap of species richness (number of distinct species-level MAGs) exclusive to the ultrasmall fraction (0–0.2μm) across oceans and depth layers: SUR (surface), DCM (deep chlorophyll maximum) and MES (mesopelagic zone). **D.** UpSet plot showing the intersection of taxa across different size fractions. The lower panel indicates which size fraction combinations are being compared, while the upper bar plot shows the number of taxa (MAGs) unique to each combination. The stacked bars are color-coded by phylum, highlighting the taxonomic composition of exclusive or shared MAGs across size classes. Oceans are indicated as AO: Arctic Ocean, NPO: North Pacific Ocean, IO: Indian Ocean, RS: Red Sea, MS: Mediterranean Sea, SAO: South Atlantic Ocean, SPO: South Pacific Ocean, SO: Southern Ocean.

Arctic and Southern Ocean samples in particular clustered separately from lower-latitude samples, reflecting the biogeographical isolation of polar bacterial communities from lower latitudes^49–52^ (Supplementary Table S2b). Notably, the ultrasmall Arctic Ocean fraction (0–0.2 µm) formed a separate cluster from all other samples, with the strongest differentiation among size fractions observed for the ultrasmall fraction (R² = 0.039–0.055, all p < 0.0001; Supplementary Table S2c). This was reinforced by taxonomic composition, where the polar oceans were dominated by Alphaproteobacteria, Gammaproteobacteria and Bacteroidota (collectively > 90%), while temperate and tropical ocean basins showed more balanced phylum distributions with contributions from Actinomycetota, Cyanobacteriota (each 5–10 %), as well as Marinisomatota. The Indian Ocean displayed the highest archaeal richness, including Nanoarchaeota (∼2.6%) and other DPANN lineages, which is consistent with their known preference for oxygen-limited or anaerobic environments (Supplementary Fig. S2, Supplementary Fig. S3)^53^.

While overall microbial diversity was reduced in the polar regions, we observed a divergence from classical latitudinal diversity patterns^52,54^ in the ultrasmall size fraction. Polar ultrasmall communities showed substantially higher species level Shannon diversity than larger size fractions (Arctic ΔH′ = 1.67; Southern Ocean ΔH′ = 2.10) compared to more modest differences in non-polar oceans (mean ΔH′ = 0.59 ± 0.35) (Supplementary Fig. S3). Notably, the Arctic ultrasmall fraction displayed one of the highest species-level diversities across all ocean basins (H′ = 3.04 ± 0.68 vs. 2.55 ± 0.65).

When focusing on size fraction niches, three main groups emerged (Fig. 2d). Out of the 1,152 non redundant MAGs, a total of 160 MAGs were confined to size fractions < 5 µm, dominated by DPANN and Patescibacterial lineages, consistent with their ultrasmall cell sizes and symbiotic or parasitic lifestyles. Conversely, 166 MAGs appeared across all size fractions, suggesting an ecologically versatile group possibly shifting between free-living, particle-associated and even larger-host or eukaryote associated lifestyles. Remarkably, 494 species were found exclusively in the ultrasmall fraction (0–0.2 µm), representing 66.8% of all undetermined bacterial species. This ultrasmall exclusive group was dominated by Pseudomonadota, Bacteroidota, and Verrucomicrobiota, and was primarily distributed in surface waters of the Arctic Ocean (Fig. 2c). This suggests the presence of a largely unexplored ecological niche of ultrasmall bacteria in polar surface waters, potentially reflecting adaptations to low temperature and nutrient scarcity.

#### Metabolic potential of the ultrasmall size fraction MAGs

To assess the metabolic potential of the 1,152 species-level MAGs from the <0.2 µm size fraction, we functionally annotated the genomes using KEGG Orthology (KO) assignments and evaluated completeness for key pathways with KEGG Decoder^55^ (Fig. 3, Supplementary Fig. S4, Supplementary file 3).

**Figure 3.**
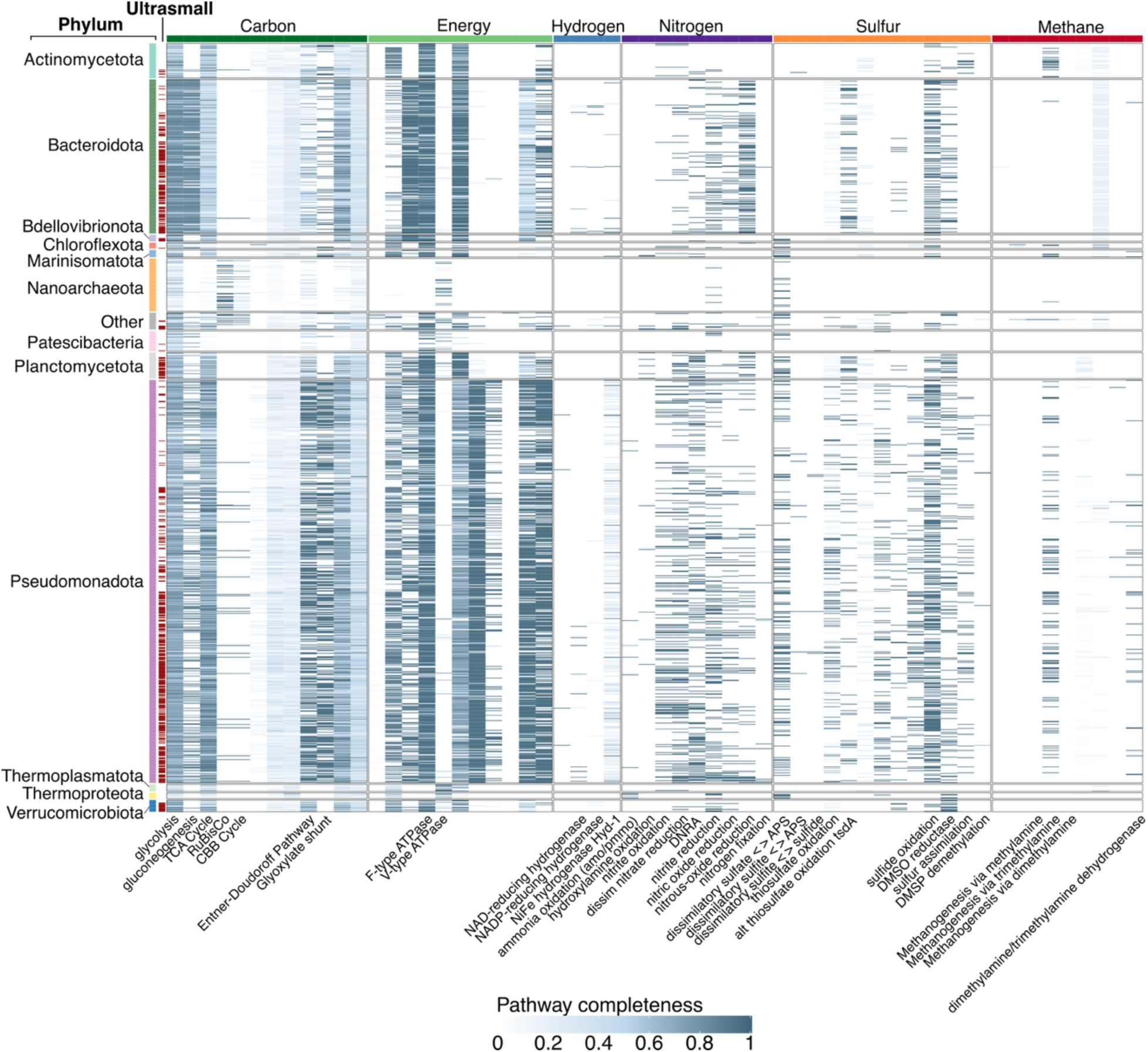
MAGs grouped by phylum. Heatmap showing pathway completeness estimated by the KEGG Decoder. MAGs (rows) are grouped and colored by phylum, with red bars indicating MAGs exclusively detected in the ultrasmall fraction. Columns represent metabolic pathways, grouped by functional categories. Blue color intensity in the heatmap reflects pathway completeness, from white (0%) to dark blue (100%). The same figure annotated with additional pathways is shown in Supplementary Fig. S4.

DPANN and Patescibacteria lineages displayed, as expected, highly reduced metabolic potential with the near-complete absence of pathways involved in major biogeochemical processes and core biosynthesis functions. Despite lacking the canonical respiratory processes, 25% of Patescibacteria MAGs encoded an F-type ATP synthase, which is hypothesized to function in reverse, contributing to ion transport or pH homeostasis rather than ATP synthesis^8,53^. Similarly, V-type ATPases occurred in all archaeal Undinarchaeia and Nanoarchaeota (2.5%) likely serving analogous proton-pumping roles.

Notably, 32.1% of Nanoarchaeota (26/81), as well as all recovered Aenigmatarchaeota and Undinarchaeota species, encoded non-canonical RuBisCO forms (II-b, III-like, II/III, IV, or IV-like) (Supplementary Fig.S5, Supplementary Data 5). Forms III and II/III function in an AMP-based CO₂-incorporating pathway rather than the CBB cycle, while form IV participates in processes such as methionine salvage and sulfur metabolism^10,16,56–58^. The prevalence of these alternative RuBisCO forms over canonical carbon fixation enzymes is consistent with the host-dependent lifestyles of DPANN archaea. Canonical forms I and II RuBisCO, which function in the Calvin–Benson–Bassham (CBB) cycle, were detected as expected in Cyanobacteria and Pseudomonadota, with unexpected occurrences in Actinomycetota (Ilumatobacter) and in one Bacteroidota MAG (Flagellimonas), a lineage not previously reported to encode RuBisCO (Supplementary Fig.S5, Supplementary Data 6, Supplementary Text).

Central carbon metabolism was generally limited, with glycolysis, gluconeogenesis, and the TCA cycle sporadically detected in Bacteroidota, Planctomycetota, Pseudomonadota, Actinomycetota, and Verrucomicrobiota. The Entner–Doudoroff pathway was comparatively widespread, particularly within Pseudomonadota (51.7%), as well as the glyoxylate shunt.

Nitrogen cycling encompassed nitrification, denitrification and potential for nitrogen fixation. Both ammonia-oxidizing bacteria (Gammaproteobacteria) and archaea (Thermoproteota, genus *Nitrosopelagicus*) were identified, with nitrite oxidation most common in Pseudomonadota (∼20% of MAGs) consistent with their role as nitrite oxidizers. Dissimilatory nitrate reduction, DNRA and denitrification pathways were broadly distributed particularly in Pseudomonadota, Bacteroidota, and Planctomycetota (Supplementary text). Additionally, 10 MAGs encoded nitrogenase genes (*nifHDK*), indicating potential nitrogen fixation capacity (see below).

Sulfur metabolism was dominated by oxidative and assimilatory processes, with widespread genes for the initial step of dissimilatory sulfate reduction but limited capacity for complete reduction. In contrast sulfide and thiosulfate oxidation pathways were common, especially in Proteobacteria, Bacteroidota, Campylobacterota, and Myxococcota. Assimilatory sulfate reduction was enriched in Cyanobacteria, Thermoproteota and Verrucomicrobiota while genes for DMSP and DMSO metabolism were detected primarily in Actinomycetota and Pseudomonadota.

Methane metabolism was largely restricted to bacterial methylotrophy, with no evidence of archaeal methanogenesis. Trimethylamine and dimethylamine utilization genes were prevalent in Pseudomonadota, Actinomycetota, and Marinisomatota, and rare occurrences in Nanoarchaeota and Bacteroidota. Methanol oxidation potential was detected exclusively in Methylophagaceae.

Hydrogen metabolism was largely absent across the ultrasmall fraction. The greatest diversity occurred in Pseudomonadota, which encoded several hydrogenase types linked to aerobic or anaerobic hydrogen oxidation, suggesting occasional mixotrophy or facultative chemolithotrophy. Scattered hydrogenases were also detected in Bacteroidota, and single representatives of Campylobacterota and Marinisomatota. Altogether, these results indicate that the ultrasmall fraction supports broad and functionally diverse metabolic potential with potential implications for global biogeochemical cycles.

### Non-cyanobacterial diazotrophs in the ultrasmall size fraction

#### Characterization of potential diazotroph MAGs from ultrasmall size fractions

While initial annotation identified nitrogenase genes (*nifHDK*) in 10 MAGs, we conducted a more comprehensive screen of all 4,058 MAGs for diazotrophy using three complementary HMM-based approaches expected to increase gene identification sensitivity^3,59,60^. This identified 474 bacterial MAGs with the catalytic *nifH* gene, with only 30 MAGs from 21 samples possessing both catalytic (*nifH, nifD, nifK*) and biosynthetic (*nifE, nifN, nifB*) genes required for functional nitrogen fixation, reinforcing that *nifH* alone is insufficient to infer diazotrophy. Many *nifH*-only hits (n=458) barely exceeded our HMM significance thresholds, further suggesting that stricter cutoffs and inclusion of *nifD/nifK* provide more reliable markers of nitrogen fixation potential.

The 30 diazotrophic MAGs originated primarily from *Tara* Oceans/Polar Circle (TOPC) (n=29) samples, with one additional MAG recovered from sponge-associated communities in the White Sea (RUSA22-1 project)^61^. Taxonomically, they belong to four bacterial classes: Gammaproteobacteria (n=21), Alphaproteobacteria (n=5), Campylobacteria (n=3) and Bacteriodota (n=1) spanning 13 genera including *Oceanobacter* (n=7), *Motiliproteus* (n=6) and *Novosphingobium* (n=3) (Fig. 4a, Supplementary text). At 95% ANI, the MAGs resolved into 13 species-level clusters, seven of which were assigned species names by GTDB taxonomy (Supplementary Table S4). Most MAGs originate from Arctic samples (Fig. 4b), with notable exceptions from the Mediterranean Sea, Atlantic, Indian and Pacific Oceans. As previously described, we observed no consistent trends towards genome reduction (Fig. 4c).

**Figure 4.**
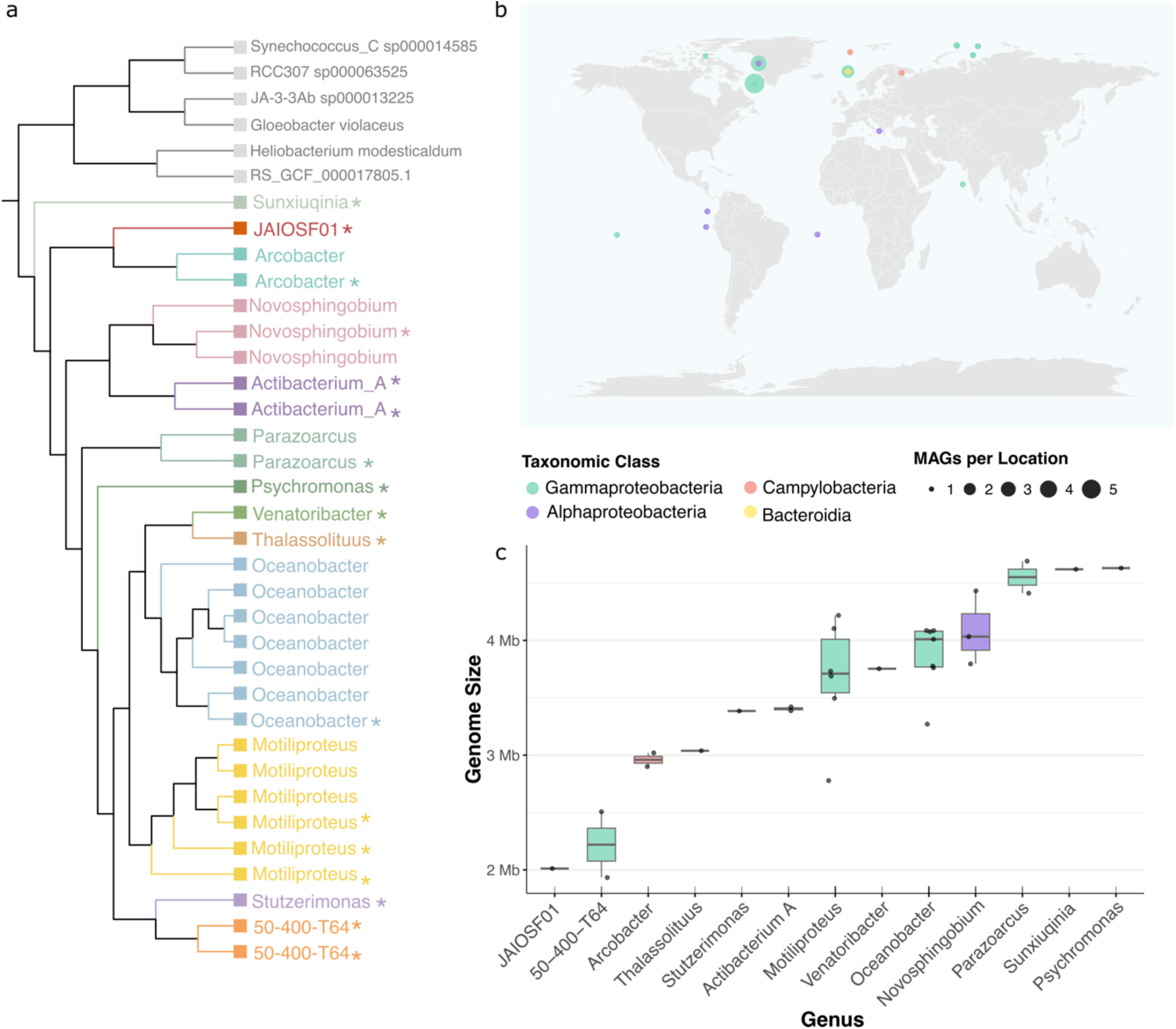
Overview of 30 diazotrophic MAGs analyzed in this study. **a**. Phylogenomic tree based on conserved marker genes constructed using PhyloPhlAn. MAGs are colored and named by their assigned genus. Outgroups are displayed in grey. Cluster representatives at strain level (99% ANI) are marked by *. **b.** Geographic origin of each MAG. The map displays sampling locations, with point size proportional to the number of MAGs recovered at each site and colors represent the phylum-level taxonomic affiliation of MAGs. **c.** Estimated genome sizes (in Mbp) of the 30 MAGs, colored by Class.

Ten MAGs lacked one or more *nif* genes, likely due to genome incompleteness, as they showed significantly lower completeness than MAGs with complete operons (Wilcoxon rank-sum test, p=0.006) (Supplementary Fig. S6). These MAGs were retained as putative diazotrophs if they clustered (at 95% ANI) with other members possessing the full set (Supplementary Table S4, Supplementary Fig. S6). Three exceptions were *Thalassolituus*, *Sunxiuqinia* (both lacking the *nifN* gene) and *JAIOSF01* (lacking *nifD*), each represented by a single MAG, which were retained with varying confidence based on taxonomic context and gene synteny (genomic neighborhood) (Supplementary text). Additionally, seven out of the thirteen diazotrophs matched previously characterized MAGs (>98% ANI) (Supplementary text, Supplementary Table S4). Three *nifH* sequences (the two *50-400-T64* strains and *JAIOSF01)* were considered novel when compared to existing *nifH* catalogs^31,36^. Only 6 of the 29 nifH sequences matched universal nifH primers, with the main mismatches occurring at *nifH4*. This is consistent with observations by Delmont et al. 2022 regarding the incompatibility of this primer with most heterotrophic bacterial diazotrophs (HBDs), a subset of NCDs lacking oxygenic photosynthesis (Supplementary Table S4).

To explore nitrogenase evolution, we placed complete concatenated *nif/vnfHDK* sequences from 28 diazotrophic MAGs in a reference phylogeny of 1,099 concatenated *nif/vnf/anfHDK* proteins^62^ (Supplementary Fig. S7). All MAGs clustered within Group I nitrogenases, except *Sunxiuqina*, which grouped with Bacteroidota in Group II, diverging from its previous *nifH*-based assignment to Group III ^32^. This discrepancy highlights the increased phylogenetic resolution provided by concatenated *nifHDK* operons, relative to single-gene trees, which are more prone to noise and artefacts derived from horizontal gene transfer. Notably, group II nitrogenases are characterized by a ∼ 50-residue conserved insertion in the *nifD* sequence^63^, which likely drives *Sunxiuqinia*’s placement. Genus-level placements of our MAGs in the *nifHDK* phylogeny aligned well with taxonomic expectations, with all MAGs forming cohesive clades with reference sequences from their respective taxa.

#### Distribution of potential diazotroph MAGs from the ultrasmall size fractions

To assess the biogeography of the identified diazotrophs, we analyzed their distribution across 210 *Tara* Oceans stations, spanning multiple depth layers, and size fractions (Fig. 5, Supplementary Fig S8). Similarly to Shiozaki et al. 2023, we observed two major distribution patterns: (i) cosmopolitan taxa with moderate relative abundances across oceans, depths, and size fractions, and (ii) Arctic-restricted taxa confined to ultrasmall fractions (<0.2 μm) with markedly elevated abundances. (Fig. 5).

**Figure 5.**
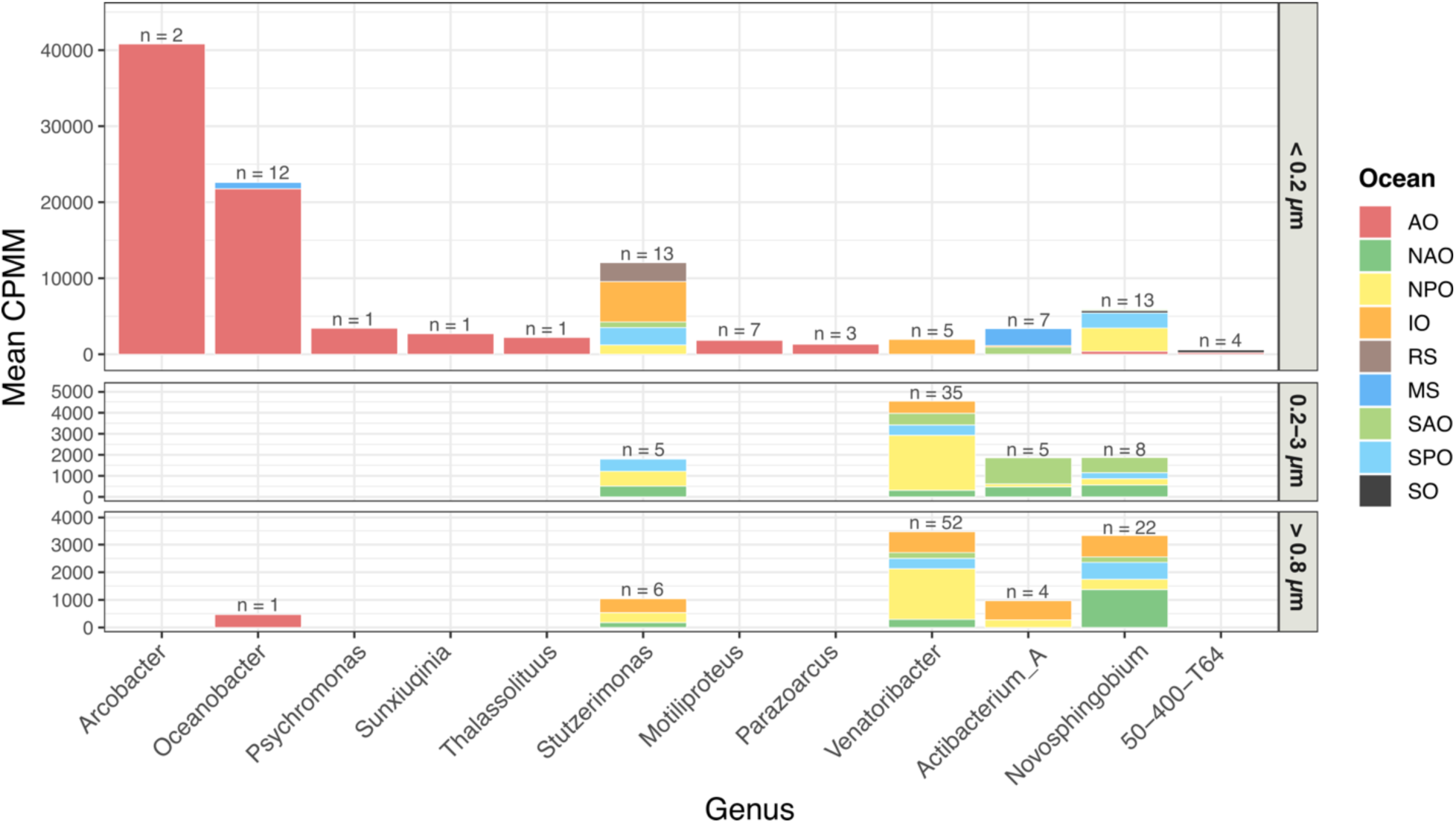
Relative abundance and distribution of potential diazotrophs from ultrasmall size fractions. MAG presence was defined as ≥60% genome coverage by reads, with abundance estimated in counts per million mapped reads (CPMM). Bars represent the mean relative abundance per genus, stacked by oceanic region. Colors indicate oceanic region: MS (Mediterranean Sea), IO (Indian Ocean), SAO (South Atlantic Ocean), SO (Southern Ocean), SPO (South Pacific Ocean), NPO (North Pacific Ocean), NAO (North Atlantic Ocean), and AO (Arctic Ocean). Distribution is shown across three size fractions: <0.2 µm, 0.2–3 µm, and >0.8 µm.

**Figure 6.**
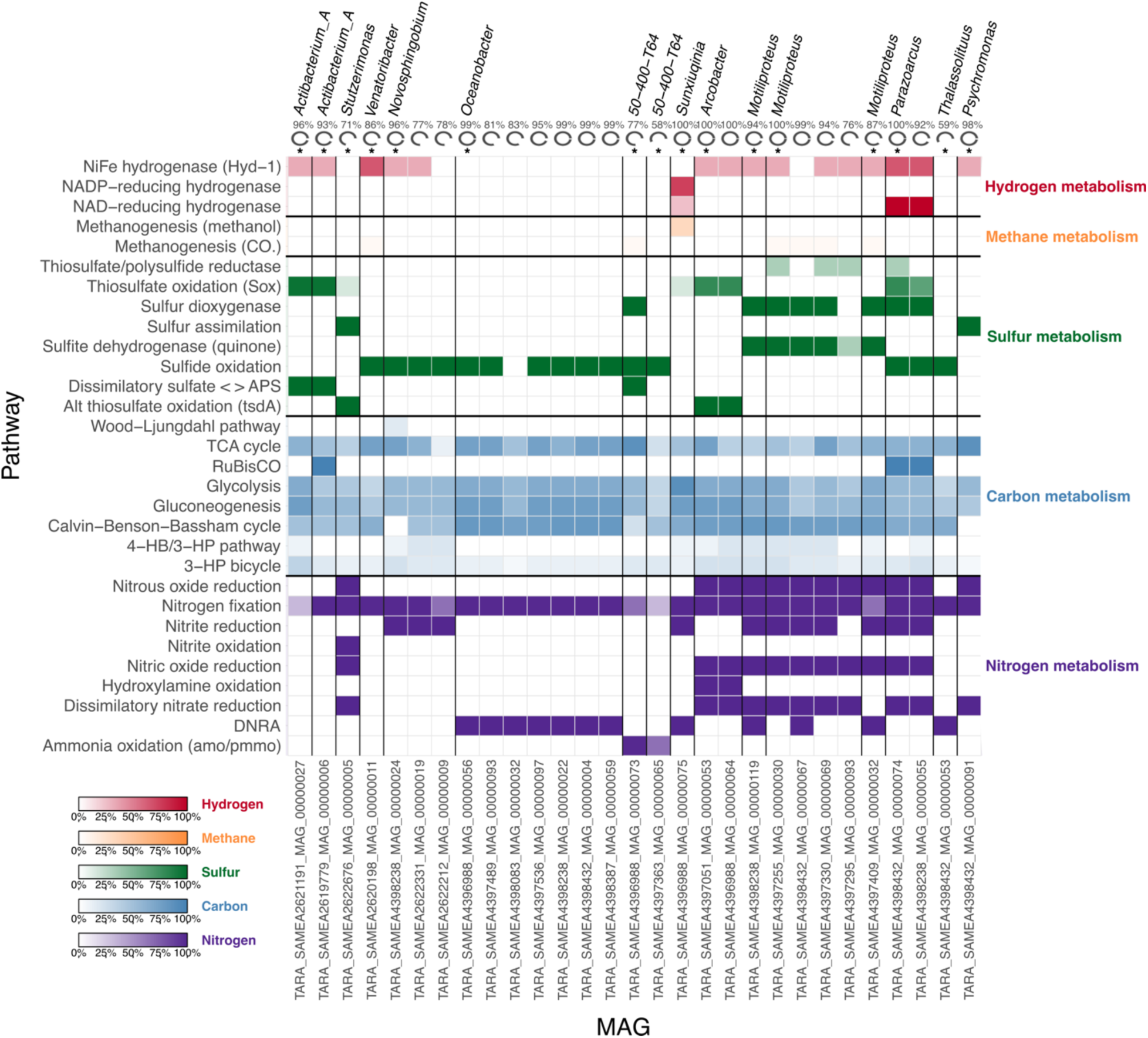
Metabolic potential of all diazotroph MAGs from ultrasmall size fractions (<0.2 µm). Key metabolic functions related to hydrogen, methane, sulfur, carbon, and nitrogen cycling of diazotroph MAGs from the ultrasmall size fraction. MAGs are grouped by genus and 99% ANI clusters are separated by black lines. Pathway completeness was determined using KEGG Decoder based on KO annotations. At the top, MAG completeness (%) is shown, with an asterisk (*) marking cluster representatives. On the right, pathways are grouped by metabolic category: hydrogen (red), methane (orange), sulfur (green), carbon (blue), and nitrogen (purple). Color intensity indicates the completeness of each KEGG pathway. The same figure annotated with pathway completeness scores is shown in Supplementary Fig. S9.

Four genera displayed cosmopolitan distributions with moderate mean relative abundances. *Venatoribacter* occurred across most major non-polar ocean basins, primarily in size fractions >0.2 μm (0.01-1.79 % relative abundance), with highest relative abundances in the North Pacific mesopelagic. *Novosphingobium* exhibited the broadest distribution, spanning nearly all ocean basins, depths, and size fractions at low relative abundances (0.01–0.26%). This aligns with previous reports of this genus across Atlantic, Indian, and Pacific basins ^3^, as well as in deep-sea environments^34^. In our dataset, it also occurred in both polar regions, but exclusively in ultrasmall fractions (0.09% Arctic, 0.04% Southern Ocean). *Stutzerimonas* and *Actibacterium_A* showed more restricted distributions, with peak abundances in the surface ultrasmall fractions of the Indian Ocean and Red Sea (0.8-1.3%) and Mediterranean Sea (0.32%), respectively.

In contrast, seven NCDs occurred exclusively or predominantly in the Arctic ultrasmall fractions, exhibiting on average six-fold higher mean normalized abundance (CPMM) than cosmopolitan taxa (Supplementary Fig. S9). This Arctic enrichment is particularly significant given that polar nitrogen fixation has only recently been recognised^26,64,65^, with summer nitrogen fixation across Arctic shelves potentially contributing 3.5 ± 0.7 Tg N y⁻¹ (∼2.7% of global BNF)^66^. Potentially ultrasmall Arctic diazotrophs were previously shown to comprise up to 1.28% of total microbial communities^32^, and 10% of ultrasmall bacterioplankton^36^. Our abundance estimates extend these findings. The Davis Strait in particular emerged as a hotspot, where *Oceanobacter* dominated at 16.7% in the mesopelagic, and co-occurred with *Motiliproteus* (0.8%), *Parazoarcus* (0.34%), *Psychromonas* (0.34%) and *Thalassolituus* (0.22%). Other Arctic-restricted ultrasmall NCDs included *Arcobacter*, reaching some of the highest relative abundances (6.7% in Norwegian Sea DCM, 1.4% at surface), while *50-400-T64* and *Sunxiuqinia* each reached 0.27% in high-latitude DCM samples. Our observations align with those of Shiozaki et al. (2023) for most overlapping taxa, with the exception of Sunxiuqinia which we detected exclusively in the ultrasmall fractions whereas they reported it across broader size classes. This discrepancy may be due to our 60% coverage threshold for MAG detection, or *Sunxiuqinia’s* filamentous morphology influencing size-fractionation behaviour.

While most ultrasmall diazotrophs were restricted to the Arctic, *50-400-T64* occurred in both the Arctic (0.27%) and Southern Ocean (0.04%) DCM ultrasmall fractions, alongside *Novosphingobium*, supporting emerging evidence for diazotrophy in Antarctic waters^27,67^. The MAG *JAIOSF01*, which was recovered from a sponge-associated microbial community in the White Sea, was absent from *Tara* Oceans, likely reflecting its coastal origin versus the open-ocean focus of these latter metagenomes.

The high abundances of diazotrophs in Arctic ultrasmall fractions indicate that standard filtration protocols (≥0.2 μm) may systematically overlook these numerically dominant populations in polar nitrogen fixation studies.

#### Metabolic potential of potential diazotrophs from the ultrasmall size fraction

The metabolic analysis of our 30 diazotrophic MAGs revealed substantial functional variation not only across taxa but also within strain-level clusters (ANI >99%). While this intra-strain variation could reflect genuine ecological diversification or incomplete MAG assembly, it raises the question if standard dereplication approaches that retain only cluster representatives may overlook functionally important genes. A pangenome-like framework retaining genes from all cluster members would better capture metabolic potential, as seen in *Actibacterium_A*, where the species representative possessed only the RuBisCO short-chain (K01602) while a closely related cluster member encoded both chains (K01601 and K01602).

Beyond nitrogen fixation, the diazotrophs displayed a wide range of nitrogen cycling capabilities that varied between cosmopolitan and Arctic taxa. Cosmopolitan genera generally showed limited nitrogen metabolism, with the exception of *Stutzerimonas,* which encoded near-complete denitrification pathways, as well as *Novosphingobium,* with nitrite oxide reduction potential. In contrast, the Arctic ultrasmall communities exhibited much greater metabolic diversity. Multiple genera (*Motiliproteus*, *Arcobacter* and *Psychromonas)* encoded near-complete denitrification, including nitrite, nitric oxide, and nitrous oxide reduction, while others (*Oceanobacter*, *Thalassolituus*, *Sunxiuqinia* and some *Motiliproteus)* possessed DNRA pathways that retains nitrogen within the ecosystem rather than releasing it through denitrification. Interestingly, *50-400-T64* encoded genes for ammonia oxidation, representing a rare nitrifier-diazotroph combination that challenges traditional views of metabolic incompatibility due to the oxygen sensitivity of the nitrogenase. *Arcobacter* also possessed the genes for hydroxylamine oxidation.

Sulfur metabolism was similarly widespread but showed distinct patterns. Cosmopolitan taxa displayed more specialized capabilities: *Stuzterimonas* encoded both sulfur assimilation and alternative thiosulfate oxidation (tsdA), *Actibacterium_A* carried the Sox pathway for thiosulfate oxidation and genes for dissimilatory sulfate reduction to APS, while *Venatoribacter* and *Novosphingobium* both showed sulfide oxidation potential.

Arctic ultrasmall communities displayed broader sulfur processing potential, with multiple taxa encoding sulfur dioxygenase (*Motiliproteus*, *Parazoarcus* and *50-400-T64),* and sulfide oxidation (*Oceanobacter, Parazoarcus, 50-400-T64, and Thalassolituus).* Additionally *Motiliproteus* encoded the sulfite dehydrogenase (quinone) and *Psychromonas* sulfur assimilation pathways.

Several diazotrophs also displayed chemolithotrophic potential. Type I RuBisCO (KEGGs K01601 and K01602) was detected in *Parazoarcus* and *Actibacterium_A,* while NAD-reducing hydrogenase in *Parazoarcus* and *Sunxiuqina* suggests potential for hydrogen-based lithotrophy. *Arcobacter* additionally contained genes for hydroxylamine oxidation The broad metabolic versatility of these ultrasmall diazotrophs, particularly within Arctic communities, suggests they may play more significant roles in marine biogeochemical cycling than previously recognized.

## Conclusion

We provide the first genome-resolved analysis of the global ocean’s ultrasmall size fraction, recovering 4,058 MAGs representing 1,152 species, over half of which are novel. The dataset includes known ultrasmall lineages (DPANN, Patescibacteria), as well as prokaryotes typically associated with larger size fractions. Over 490 species were found exclusively in the ultrasmall size fraction, predominantly in Arctic waters, suggesting a previously unknown ultrasmall polar-adapted niche. Among these we identified 13 putative non-cyanobacterial diazotrophs through complete nitrogenase operon screening, with an ultrasmall restricted polar community, displaying elevated abundances in particular in the Davis Strait.

These MAGs display remarkable metabolic diversity, spanning nitrate reduction, DNRA, sulfate reduction and methylotrophy. This functional diversity may indicate a community with complementary metabolic strategies or alternatively an assemblage of populations in reduced physiological states that may not be actively thriving in this niche. The mechanisms underlying their presence in the <0.2 μm size fraction still remain unclear and need further investigation. Possibilities include genuinely ultrasmall cell sizes, dormancy or starvation-induced size reduction, or selective enrichment of rare taxa during size fractionation. Although repeated detection across independent samples partially mitigates contamination concerns, we cannot entirely exclude artifacts from size fractionation.

Moreover, to discriminate between active metabolic strategies and reduced physiological states, as well as quantify biogeochemical impacts, metabolic rate measurements with cell abundance estimates are essential. In parallel, long-read sequencing will be necessary to improve MAG completeness and refine functional interpretation.

Despite these uncertainties our findings demonstrate that the ultrasmall fraction represents an overlooked reservoir of microbial diversity with potential roles in key biogeochemical processes, in particular in the Arctic. These results highlight the necessity of including the ultrasmall size fraction in future microbial diversity surveys as well as taking it into consideration in biogeochemical models to achieve a more complete understanding of ocean microbial systems.

## Materials and Methods

### Reconstruction of MAGs in ultrasmall size fraction samples across the global ocean

The Ocean Microbiomics Database (OMDB; omdb.microbiomics.io) compiles 209 publicly accessible studies from key publications in marine metagenomics. For each study, the raw sequencing reads were retrieved from the European Nucleotide Archive (ENA) and processed to reconstruct MAGs following a benchmarked workflow described in Paoli et al. (2022)^4^. Only medium- and high-quality MAGs were retained, using thresholds of ≥50% completeness and ≤10% contamination and ≥90% completeness and ≤5% contamination, respectively. For each metagenomic sample in OMDB, metadata were retrieved from ENA by their biosample identifier. Information on size fractions was manually curated based on information available in the respective original publications. In total, a lower and upper size fraction was assigned to 9,742 out of 12,347 samples. Samples with the size fractions ‘0-0d1’, ‘0-0d2’ and ‘0d1-0d2’, were included in this study, which contained a total of 4,058 MAGs. All MAGs analyzed in this study (Supplementary Dataset S1) are publicly accessible at: omdb.microbiomics.io.

Ultrasmall size fraction samples are commonly processed using FeCl₃ precipitation to remove viral particles prior to filtration, enriching for prokaryotic cells in this size class. In this protocol, <0.2 µm water is complemented with FeCl₃ to precipitate viral particles for 1 to 24 hours, then the resulting precipitate is filtered on a 0.8 µm PC filter and stored at +4°C. This preprocessing step should be considered when interpreting the composition of these metagenomic datasets.

### Taxonomic assignation

Genome taxonomic classification was performed using GTDB-Tk v2.4.0^68^ with GTDB reference data release r220. The classify_wf workflow was run with default parameters.

As part of the pipeline, the following tools were used: Mash v2.3^69^, Skani v0.2.2^70^, Prodigal v2.6.3^71^, HMMER v3.4^72^, and pplacer v1.1.alpha19^73^.

### Genome redundancy analysis

Pairwise average nucleotide identity (ANI) was calculated using skani (v0.2.2)^70^ with default parameters for the 4,058 genomes within the OMDB small size fraction samples. MAGs with >95% or >99% ANI were considered redundant at the species and strain levels, respectively. The longest MAG (by assembly size) within each cluster was selected as the representative genome. This resulted in a non-redundant MAG dataset of 1,152 species clusters and 1,824 strain clusters.

### Biogeography of MAGs

To assess the distribution of dereplicated MAGs, all *Tara* Oceans metagenomes (all stations and size fractions) were mapped to the species-level MAG dataset as described in [Carradec, Pelletier, et al. 2018]^74^. Briefly, raw reads were mapped using bwa-mem (version 2.2.1)^75^ with a 95% identity threshold, under the “random best match” mode. Duplicates were removed with the samtools^76^ suite, and matches covering at least 70% of the length of the read were filtered using bamFilters. MAGs were considered present in a sample if ≥60% of their genome length was covered by reads.

Genome abundance was calculated as counts per million microbial genomes (CPMM), a length-normalized metric adapted from Shiozaki et al.^32^ that is conceptually similar to TPM used in transcriptomics. CPMM was calculated as follows:

i. length-normalized counts per genome (Gi) were computed as Gi = 1000 × Ci / Li, where Ci is the number of mapped reads on the genome i and Li is the genome i size.
ii. these values were then normalized to counts per million targets (CPMT = 10⁶ × Gi / ΣGi) across all MAGs within each sample.
iii. CPMM was calculated by scaling CPMT by the proportion of reads mapped to the MAG set relative to total reads in the sample: CPMM = CPMT × (total_mapped_reads_on_all_genomes / total_raw_reads).

CPMM reflects the MAG abundance within the total microbial community; for example, CPMM = 1000 corresponds to 0.1% abundance.

For comparative analyses across size fractions, the original size fractions from Tara Oceans samples were grouped into four broad categories: ultrasmall (0–0.2 µm), free-living (0.2–5 µm), particle-attached (0.8–20 µm), and large particles/aggregates (0.8–∞ µm). The ultrasmall fraction combined samples labeled as 0–0.2 µm and 0.1–0.2 µm. The free-living fraction included 0.2–0.45 µm, 0.2–1.6 µm, 0.22–3 µm, 0.45–0.8 µm, 0.8–3 µm, and 0.8–5 µm. The particle-attached fraction combined 0.8–20 µm, 1.6–20 µm, 3–20 µm, and 5–20 µm. The large particle fraction included 0.8–∞ µm, 20–180 µm, 180–2000 µm, 3–∞ µm, 20–200 µm, and 0.8–200 µm. For the diazotroph distribution analysis, the particle-attached and large particle fractions (0.8–20 µm and 0.8–∞ µm) were further collapsed into a single category. Analyses were performed across three depth layers sampled by Tara Oceans: surface (SUR), deep chlorophyll maximum (DCM), and mesopelagic (MES).

### Metabolic analysis

We inferred functional annotation of predicted proteins from our MAG contigs using anvi’o’s *anvi-run-kegg-kofams* (v7.1)^77,78^. Gene prediction was performed with Prodigal v2.6.3^71^ as part of the anvi’o workflow. KEGG Orthologs (KOs) were assigned based on the KOfam HMM database using HMMER v3.4^72^ with default parameters. Metabolic pathway completeness was estimated from these KO annotations using KEGG Decoder^55^, and pathway completeness results were visualized in R with custom scripts adapted from KEGG Decoder outputs.

### Statistical analysis

#### Community Composition Analysis

Non-metric multidimensional scaling (NMDS) ordination was performed on MAG abundance tables, using the metaMDS function from the vegan package (v2.7.1)^79^ in R. Bray-Curtis dissimilarity was used as the distance metric, with ordination computed in two dimensions (k=2) and a maximum of 100 iterations to achieve convergence. Seven samples (37MES01QQRR12, 36DCM0AACC11, 136SUR1MMQQ11, 93SUR01GGZZ11, 18SUR1SSUU11, 80SUR1QQSS11, and 137DCM0AACC11) were excluded from the analysis as they appeared as clear outliers (Supplementary Fig. S10). Permutational multivariate analysis of variance (PERMANOVA)^80^ was performed to partition variance in the Bray-Curtis dissimilarity matrix among ocean basins, size fraction categories and depth categories, using the adonis2 function from the vegan package with 10,000 permutations.

Shannon diversity (H′) was calculated from MAG relative abundances (CPMM-normalized) using the diversity() function in vegan^81^ with the “shannon” index. Diversity was computed at both the species and phylum levels after aggregating abundances accordingly. All analyses and visualizations were performed in R (v4.3.1) using tidyverse^82^ and ggplot2^83^.

#### Genome Completeness and Operon Status

Genome completeness values were compared between MAGs with complete operons (*nif*HDKENB or *vnf*H-DKENB) versus those with incomplete operons using the Wilcoxon rank-sum test with the wilcox.test() function in R.

### Identification of diazotroph MAGs

We identified putative diazotrophs using three complementary hidden Markov model (HMM) approaches. We used Pyrodigal (v3.4.1)^71,84^ to predict genes in each considered MAG, and produced a catalog of 11,417,472 coding sequences that was then screened for nitrogen fixation genes (*nifHDKENB*) using (i) TIGRFAM HMM profiles as implemented in Delmont et al. (2022)^3^, (ii) KEGG Ortholog (KO) annotations^59^, and (iii) the NFixDB HMM collection^60^, which includes profiles for *nifHDK, anfHDK, vnfHDK*, as well as related pseudogenes (*chlB, chlL, chlN, nflD*, and *nflH*). HMM searches were performed using PyHMMER (v0.10.11) with default parameters^84,85^. Then results were grouped by genome, selecting those meeting the quality and size fraction criteria described above. HMM gene hits were retained if they had a bitscore >50 or an e-value <9.9e-15, and best hits were selected between the HMMs.

MAGs were screened for nitrogenase gene sets from three systems: *nif*HDKENB (molybdenum nitrogenase - full operon), *anf*HDK (iron-only nitrogenase - partial operon), *vnf*HDK (vanadium nitrogenase - partial operon). However, as noted by Belanger et al. 2024^60^, HMMs can have difficulty distinguishing between *nifH* and *vnfH* due to their close similarity. To account for this, we combined *nifH* and *vnfH* hits in our analysis, always selecting the highest-scoring match between the two. Finally, MAGs were considered candidate diazotrophs if they possessed any of the following gene sets: *nif*HDKENB (complete molybdenum nitrogenase), *anf*HDK (partial iron-only nitrogenase), *vnf*HDK (partial vanadium nitrogenase), *vnf*H-*nif*DKENB (where *vnf*H was the top hit over *nif*H, with remaining genes from the *nif* operon). To increase confidence in gene calls, we required that *H/D/K* and *E/N* genes respectively co-occur on the same scaffold within 10 genes of one another. Additionally MAGs with incomplete nitrogenase operons were retained if they clustered at ≥95% ANI with other MAGs in the dataset that possessed the full operon. In our final dataset, no MAGs possessed the alternative nitrogenase gene sets (*anf*HDK or *vnf*HDK). All identified diazotrophs contained genes from the molybdenum nitrogenase system (*nif*), with the *nif*H/*vnf*H ambiguity handled as described above.

### Phylogenomic analysis of diazotroph MAGs

We constructed a phylogenomic tree of our putative diazotrophs using PhyloPhlAn v.3^86^, which infers genome-wide microbial phylogenies based on ∼400 conserved marker proteins. For our analysis, we applied the medium diversity marker set with default pipeline parameters.

### Phylogenetic analysis of nitrogenase genes

To examine the phylogenetic placement of functional marker genes from our MAGs, we reconstructed maximum-likelihood trees for both nitrogenase and RuBisCO sequences. For nitrogenase, we added the amino acid sequences of concatenated *nifH*, *nifD*, and *nifK* to the reference *nifHDK* database of 1,099 sequences from Pi et al. (2024)^62^. For RuBisCO, 63 large-subunit sequences (KO: K01601) identified in our MAG dataset were aligned against the RuBisCO large-chain reference profile from Jaffe et al. 2019^16^. Sequences were aligned using MUSCLE v5.1^87^, and maximum-likelihood phylogenies were inferred with IQ-TREE v2.1.4^88^ using ModelFinder^89^ for automatic model selection. In both cases, the LG+R10 model was chosen as the best-fitting model, and branch support was assessed with 1,000 ultrafast bootstrap replicates. The nitrogenase tree was rooted using concatenated outgroup sequences from Bch/ChlLNB and BchXYZ proteins, while the RuBisCO tree was left unrooted. Final trees were visualized in TreeViewer^90^.

### *NifH* sequence novelty

We assessed the sequence novelty of our 29 *nifH* genes from potential diazotrophs by comparing them against the nifH database described in Pierella et al. 2021 and the nifH catalogue published by Turk-Kubo (2023)^31^. As introduced in Delmont et al. (2022)^91^, sequences were considered novel if they shared less than 98% nucleotide identity over a minimum alignment length of 200 base pairs.

## Acknowledgments

We would like to thank Dominic Eriksson and Jonas Richter for their assistance with metadata curation, Hans Ruscheweyh for data management and computational support, and the ETH IT services and HPC facilities for granting access to the EULER high performance cluster. This article is contribution number XXX of *Tara* Oceans.

## Data Availability

All MAGs and associated metadata are available through the Ocean Microbiomics Database (OMDB) at omdb.microbiomics.io. The associated MAG and sample IDs are in the Dataset S1. The code and all supplementary datasets supporting the findings of this study (Supplementary Datasets S1–S5) will be made available upon publication.

## Author Contributions

C.A.H., S.M.V., S.S, S.G.A. and C.B designed the research. C.A.H., S.M.V. and E.P performed the research. C.A.H., S.M.V., S.S. and L.J.U analyzed the data. Everyone wrote the paper.

## Competing Interest Statement

The authors declare no competing interest.

## Funding

Work in the Bowler lab has been funded by the European Research Council (ERC) under the European Union’s Horizon 2020 research and innovation program (Diatomic; grant agreement No. 835067) and BlueRemediomics (grant agreement No. 101082304), the French Government “Investissements d’Avenir” programs MEMO LIFE (ANR-10-LABX-54), PSL Research University (ANR-11-IDEX-0001-02), the Agence Nationale de la Recherche (DIM ANR-21-CE02-0021-01), and the Fondation BNP Paribas Climate and Biodiversity Initiative. L.J.U. acknowledges funding from the Peter und Traudl Engelhorn Stiftung zur Förderung der Lebenswissenschaften,S.M.V. acknowledges funding from the Human Frontier Science Program (HFSP) through a postdoctoral fellowship [LT0050/2023-L], S.S. acknowledges funding from the Swiss National Science Foundation (grant number 205320_215395).

## Notes

### Competing Interest Statement

The authors have declared no competing interest.

