## Supplementary Information for "The ultrasmall ocean microbiome: a reservoir of microbial diversity and nitrogen fixation"

† Deceased

**Table of contents**

**Supplementary text**
**Figure S1.** Number of species-level MAGs assembled from the <0.2μm size fraction by ocean and phylum.
**Figure S2.** Stacked bar plots showing the percent of total CPMM of the 1,152 MAGs from the ultrasmall size fraction.
**Figure S3.** Shannon diversity indexes by ocean basin at A. species level B. phylum level.
**Figure S4.** Metabolic pathway completeness for 1,152 species-level ultrasmall-fraction MAGs grouped by phylum.
**Figure S5.** Phylogenetic tree of long chain RuBisCO (K01601) amino acid sequences present in <0.2 μm MAGs.
**Figure S6.** Heatmap showing nif gene presence and operon completeness for all potentially diazotrophic MAGs in the ultrasmall size fraction.
**Figure S7**. Phylogeny of concatenated nifHDK nitrogenase proteins from diazotroph MAGs from ultrasmall size fractions with respect to reference sequences.
**Figure S8.** Distribution of potential diazotrophs by depth.
**Figure S9.** Metabolic potential of all diazotroph MAGs from ultrasmall size fractions (<0.2 μm).
**Figure S10.** Non-metric multidimensional scaling (NMDS) based on Bray-Curtis dissimilarity of MAG abundance profiles.
**Table S1**. Source studies contributing MAGs to the ultrasmall size fraction dataset.
**Table S2.** PERMANOVA analysis of MAG community structure.
**Table S3.** Shannon diversity between A. ultrasmall and larger size fractions. B. Ocean basin.
**Table S4.** Summary of the 17 strain level cluster representatives (99% ANI) of potential diazotrophs. Bold MAGs with a * indicate cluster representatives at ANI 95%.
**Supplementary Datasets**

### Supporting Information Text

#### Supplementary overview of over 4,000 MAGs from the ultrasmall size fraction.

**Supplementary analysis on MAG genome size.** The largest MAG in our dataset (16.93 Mb, assigned to *SCGC-AAA160-P02* sp913060575) substantially exceeds expected genome sizes for this lineage and is likely due to a contaminated assembly. Quality assessment revealed differences between binning evaluation tools: Anvi'o estimates showed completeness of 78.87% and contamination of 7.04%, while CheckM estimates indicated completeness of 97.38% and contamination of 39.36%. The high contamination estimated from CheckM, combined with extreme fragmentation (3,104 scaffolds), strongly suggests that the MAG contains genomic material from multiple organisms. When examining the 13 other genomes of *SCGC-AAA160-P02* sp913060575 in OMDB their sizes ranged from 1.2 to 2.7 Mb, which is consistent with genome sizes for this species. In this case, selecting the cluster representative based on genome quality metrics (e.g., completeness and contamination) would likely provide a more appropriate representative.

#### Supplementary metabolic analysis of the ultrasmall size fraction MAGs

**Detailed RuBisCO diversity in ultrasmall MAGs.** A more detailed analysis of RuBisCO diversity revealed a predominance of non-canonical forms among DPANN archaea. In particular, Nanoarchaeota, Aenigmatarchaeota, and

Undinarchaeota primarily encoded RuBisCO forms III-b, III-like, II/III, IV, or IV-like, consistent with roles in AMP-based $CO_2$ incorporation or alternative metabolic processes rather than canonical carbon fixation.

Beyond expected carriers, we detected RuBisCO type I in a single Actinomycetota MAG (Ilumatobacter), and in one Bacteroidota MAG (Flagellimonas), a lineage not previously reported to encode RuBisCO. Actinomycetota are more commonly known to encode form IE RuBisCO; however, autotrophic growth via the CBB cycle has been confirmed in only a few cultured representatives[1]. Notably, our Ilumatobacter MAG lacked hydrogenase genes typically linked to form IE-mediated chemolithoautotrophy. While the absence of specific genes in metagenome-assembled genomes should be interpreted with caution due to potential incompleteness, this suggests that its RuBisCO may serve a non-autotrophic or salvage function.

**Nitrogen cycling pathway resolution.** Downstream nitrification steps, such as hydroxylamine oxidation, were rare, while nitrite oxidation was most common in Pseudomonadota (~20% of MAGs), consistent with their role as nitrite oxidizers, with sporadic detection in Chloroflexota and Hydrogenedentota.

Genes encoding dissimilatory nitrate reduction and Dissimilatory nitrate reduction to ammonium (DNRA) were widespread, and particularly enriched in Planctomycetota and Pseudomonadota, supporting their role in nitrate turnover and nitrogen retention. Denitrification pathways were broadly distributed: nitrite reduction (nirK/nirS), nitric oxide reduction (norBC), and nitrous oxide reduction (nosZ) occurred across multiple phyla, with strong representation in Pseudomonadota, Bacteroidota, and Planctomycetota.

### Supplementary characterization of potential diazotroph MAGs from ultrasmall size fractions

**Taxonomic and genomic resolution.** At 95% ANI, the 30 diazotrophic MAGs resolved into 13 species-level clusters and into 17 clusters at the strain level (99% ANI). Phylogenomic reconstruction using ~400 conserved marker genes revealed clustering patterns consistent with GTDB taxonomy. The full taxonomic breakdown includes: *Oceanobacter* (n=7), *Motiliproteus* (n=6), *Novosphingobium* (n=3), *Parazoarcus* (n=2), *50-400-T64* (n=2), *Actibacterium_A* (n=2), *Arcobacter* (n=2) and single representatives of *Psychromonas, Stutzerimonas, Sunxiuqinia, Thalassolituus, Venatoribacter* and *JAIOSF01* (Arcobacteraceae family – RUSA22-1 project).

Geographic origins outside the Arctic include two *Actibacterium* MAGs from the Mediterranean Sea and South Atlantic Ocean, *Venatoribacter* from the Indian Ocean, two *Novosphingobium* MAGs from Pacific waters, and *Stutzerimonas* from the South Pacific Ocean.

**Diazotroph potential in MAGs with incomplete operons.** The *Sunxiuqinia* MAG showed 99.83% ANI with the previously published diazotroph *Arc-Bactero*[2], and a significant second-best BLAST hit for *nifN* (bit score 268.94, e-value 4.04e-81) was identified in a gene annotated as a second *nifK* copy, consistent with *nifN* arising from *nifK* gene duplication. Based on this evidence, the *Sunxiuqinia* MAG was considered diazotrophic. *JAIOSF01*, an *Arcobacteraceae* MAG, contains *nifH* and *nifD and nifENB*, with *nifD* located at the end of a contig. As *nifHDK* typically form an operon structure, this positioning suggests that *nifK* absence is more likely due to contig fragmentation rather than true absence. The *Thalassolituus* MAG lacks *nifN* but contains a complete *nifHDK* operon; given the presence of confirmed diazotrophs within *Thalassolituus* provides taxonomic support for nitrogen fixation potential within *Thalassolituus*, this MAG was retained as a lower-confidence putative diazotroph.

**Comparison with previously published diazotrophs.** Seven out of the twelve diazotrophs matched previously characterized MAGs (>98% ANI): *Novosphingobium* (ANI = 98.49%) and *Venatoribacter* (ANI = 99.97%) from Delmont et al. 2022[3] and *Psychromonas* (ANI = 99.62%), *Oceanobacter* (ANI = 99.37%), *Motiliproteus* (ANI = 99.77%), *Arcobacter* (ANI = 99.99%) and *Sunxiuqinia* (ANI = 99.83%) from Shiozaki et al. 2023[2]. Notably, Arc-Gamma-03 (99.37% ANI with Oceanobacter), was also reported as Candidatus Thalassolituus haligoni by Rose et al. 2024[4]. Additionally, Arc-Gamma-04 was not classified as a diazotroph by Shiozaki et al. due to apparent *nifB* absence. In contrast, all except one *Motiliproteus* MAGs in our dataset contain the *nifB* gene, supporting their diazotroph classification.

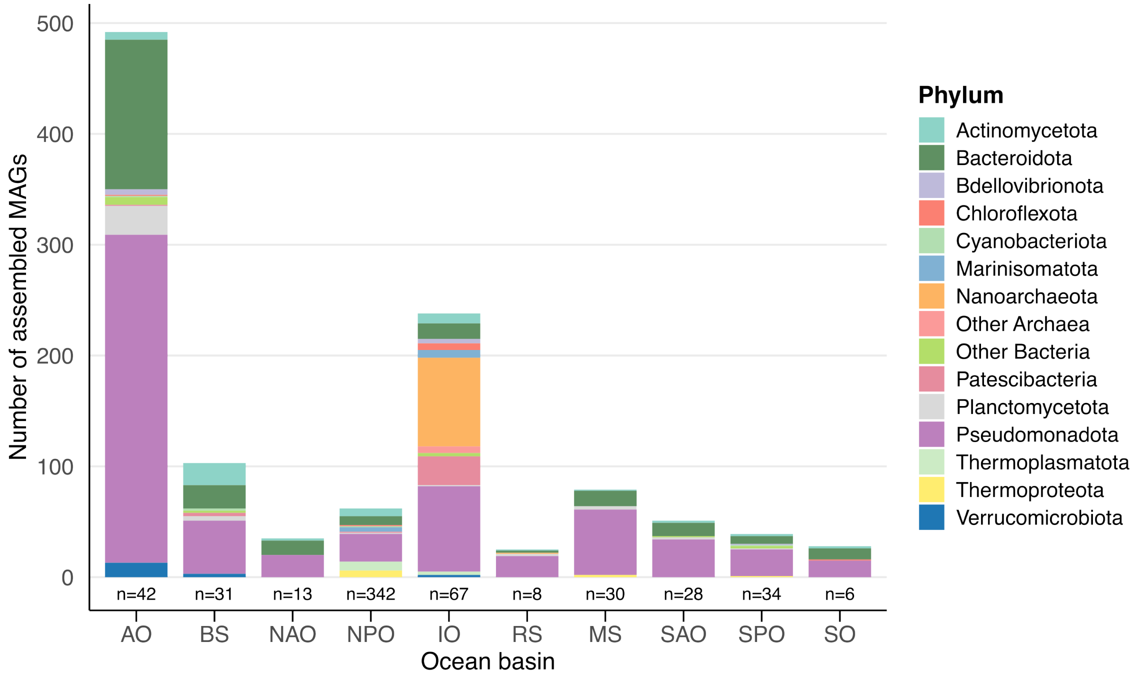


**Figure S1. Number of species-level MAGs assembled from the <0.2 µm size fraction by ocean and phylum. n indicates the number of samples.** Oceans are indicated as AO: Arctic Ocean, BS: Baltic Sea, NAO: North Atlantic Ocean, NPO: North Pacific Ocean, IO: Indian Ocean, RS: Red Sea, MS: Mediterranean Sea, SAO: South Atlantic Ocean, SPO: South Pacific Ocean, SO: Southern Ocean

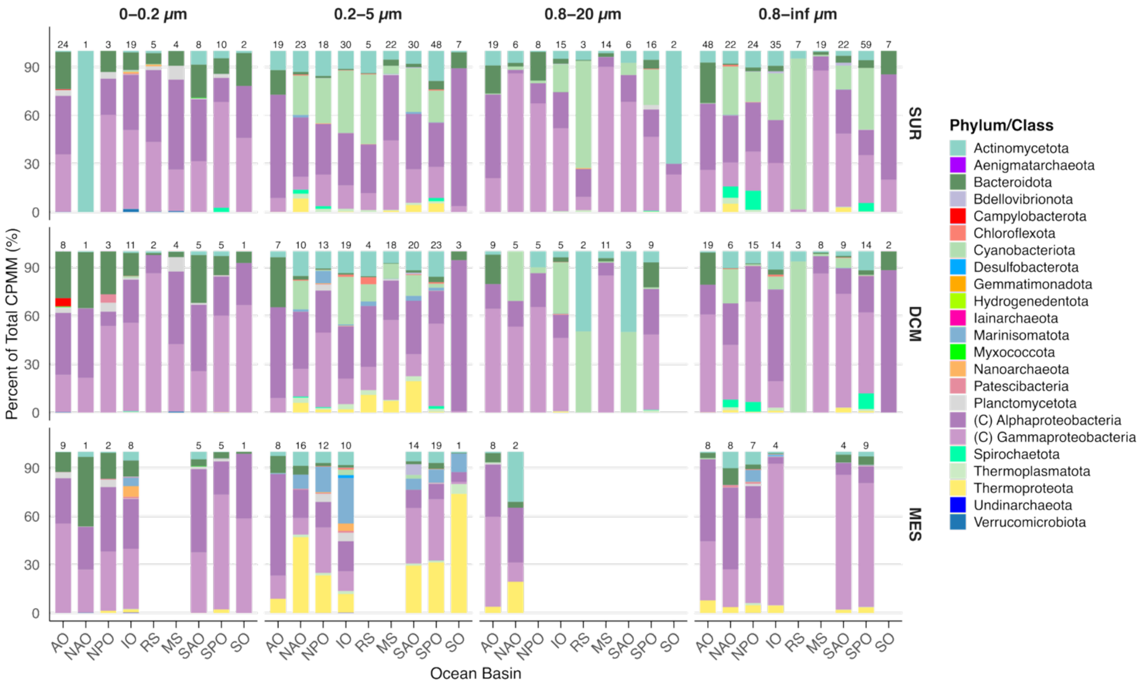


**Figure S2. Stacked bar plots showing the percent of total CPMM of the 1,152 MAGs from the ultrasmall size fraction** (phylum level, except for Pseudomonadota, where the Class (C) is shown) in each ocean basin, size fraction and depth. Numbers above the bars indicate the number of samples. Oceans are indicated as AO: Arctic Ocean, NPO: North Pacific Ocean, IO: Indian Ocean, RS: Red Sea, MS: Mediterranean Sea, SAO: South Atlantic Ocean, SPO: South Pacific Ocean, SO: Southern Ocean. Depths are indicated as SUR: Surface, DCM: Deep Chlorophyll Maximum, MES: Mesopelagic; and sorted top to bottom in different rows.

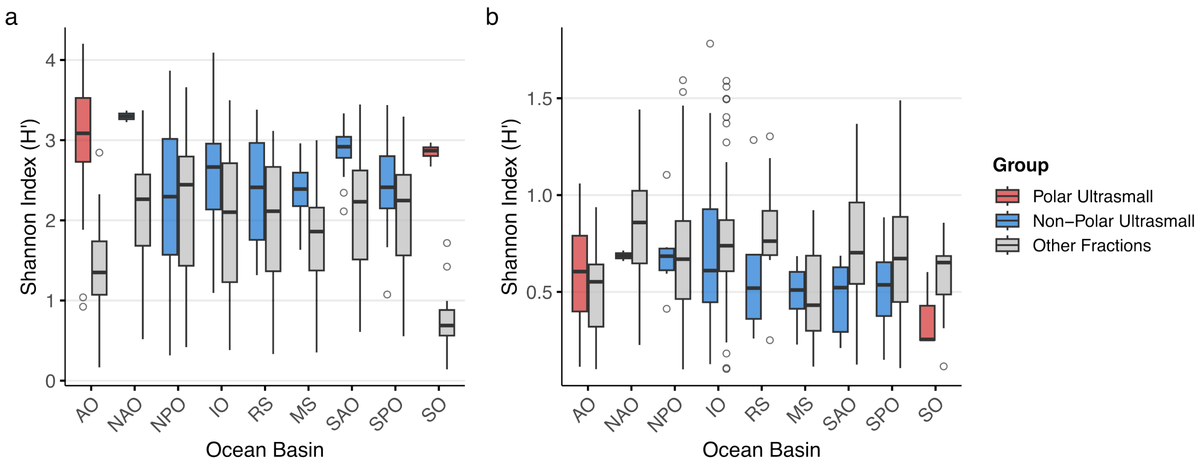


**Figure S3. Shannon diversity indexes by ocean basin at a. species level b. phylum level.** Oceans are indicated as AO: Arctic Ocean, NPO: North Pacific Ocean, IO: Indian Ocean, RS: Red Sea, MS: Mediterranean Sea, SAO: South Atlantic Ocean, SPO: South Pacific Ocean, SO: Southern Ocean.

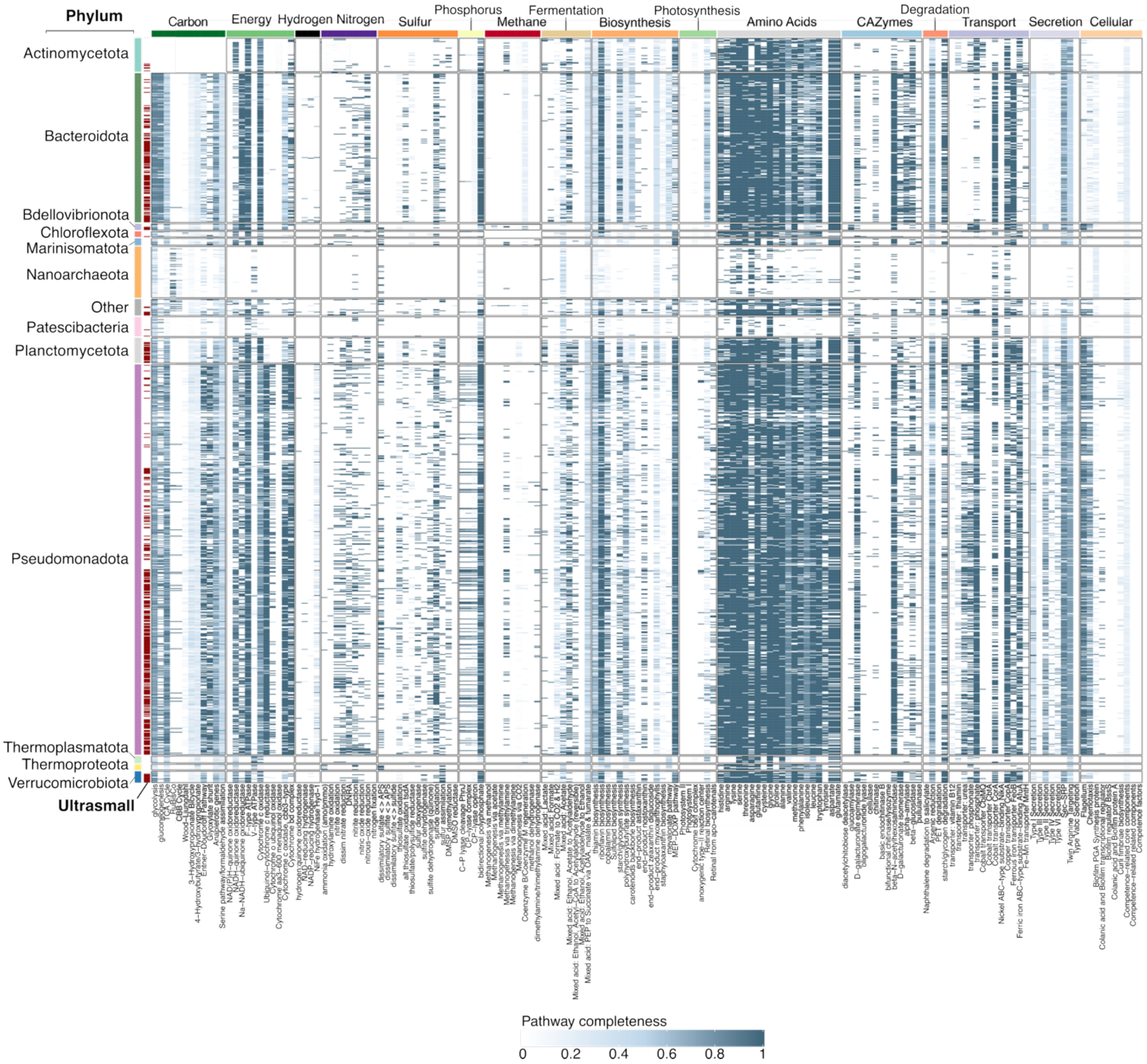


**Figure S4. Metabolic pathway completeness for 1,152 species-level ultrasmall-fraction MAGs grouped by phylum.** Heatmap showing pathway completeness estimated by the KEGG Decoder. MAGs (rows) are grouped and colored by phylum, with red bars indicating MAGs exclusively detected in the ultrasmall fraction. Columns represent metabolic pathways, grouped by functional categories. Blue color intensity reflects pathway completeness, from white (0%) to dark blue (100%).

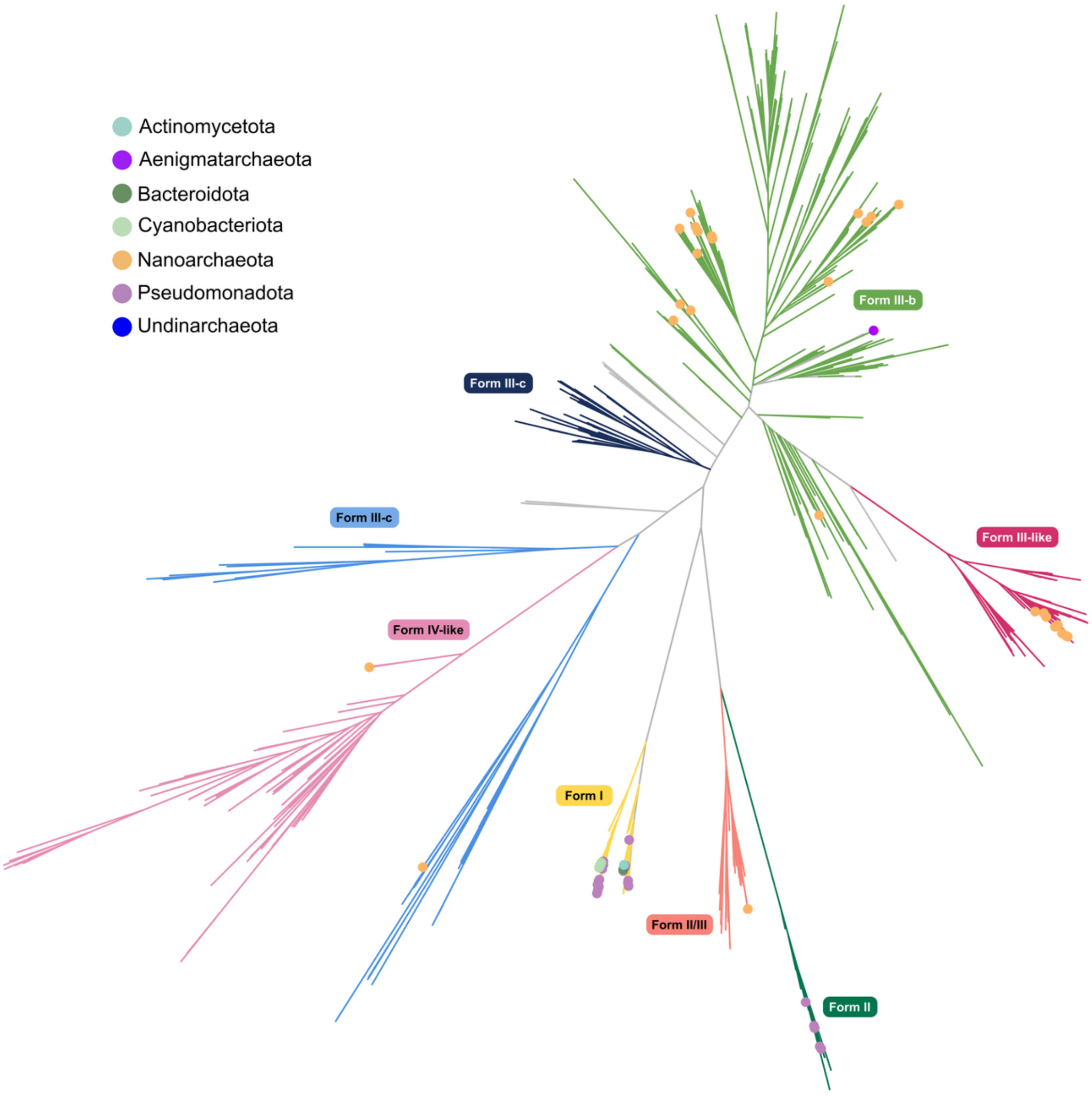


**Figure S5. Phylogenetic tree of long chain RuBisCO (K01601) amino acid sequences present in <0.2 μm MAGs.** Branches are colored according to RuBisCO form included in the colored rectangles, and terminal leaves are colored by the phylum of the <0.2 μm MAGs containing RuBisCO genes.

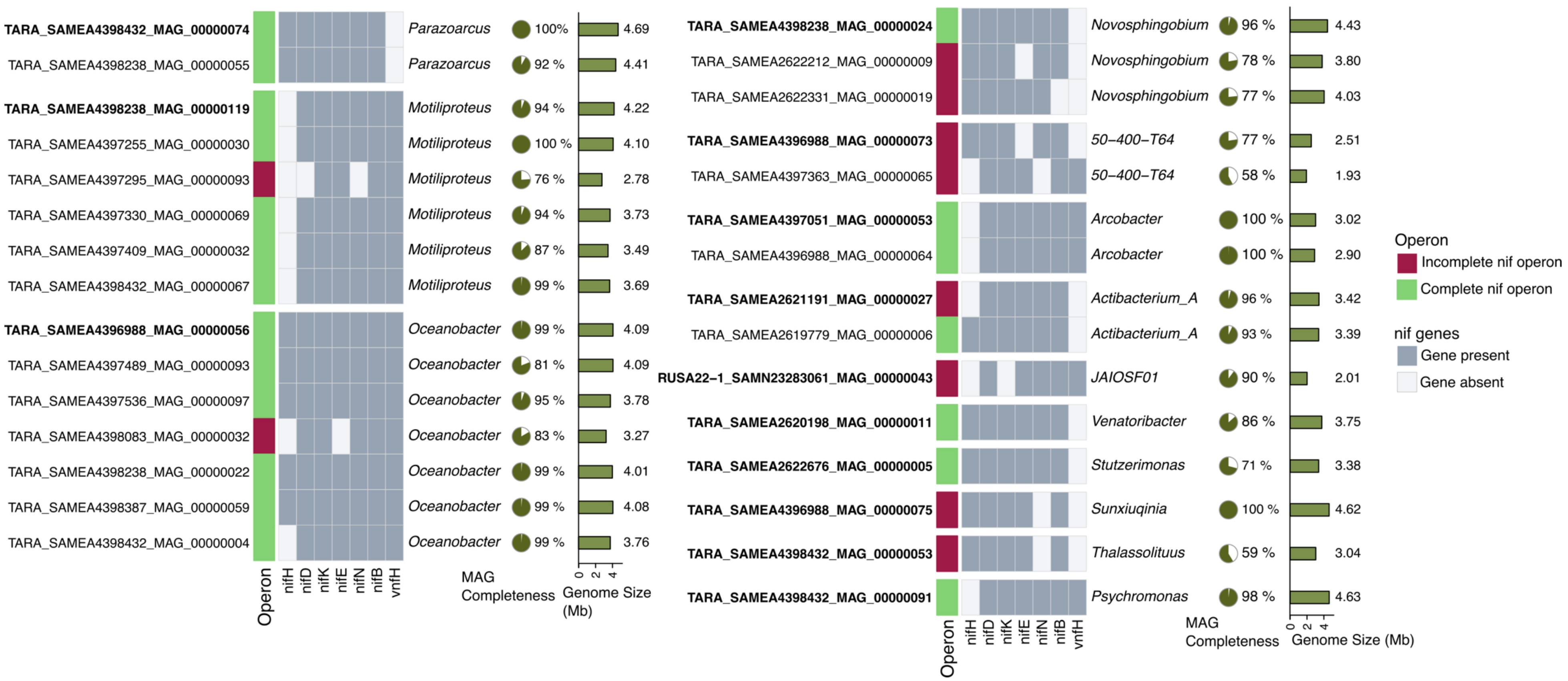


**Figure S6. Heatmap showing *nif* gene presence and operon completeness for all potentially diazotrophic MAGs in the ultrasmall size fraction.** MAGs are represented as clusters of 95% ANI with the cluster representative in bold. An operon is considered complete if it possesses *nifHDKENB* or *vnfH/nifDKENB.*

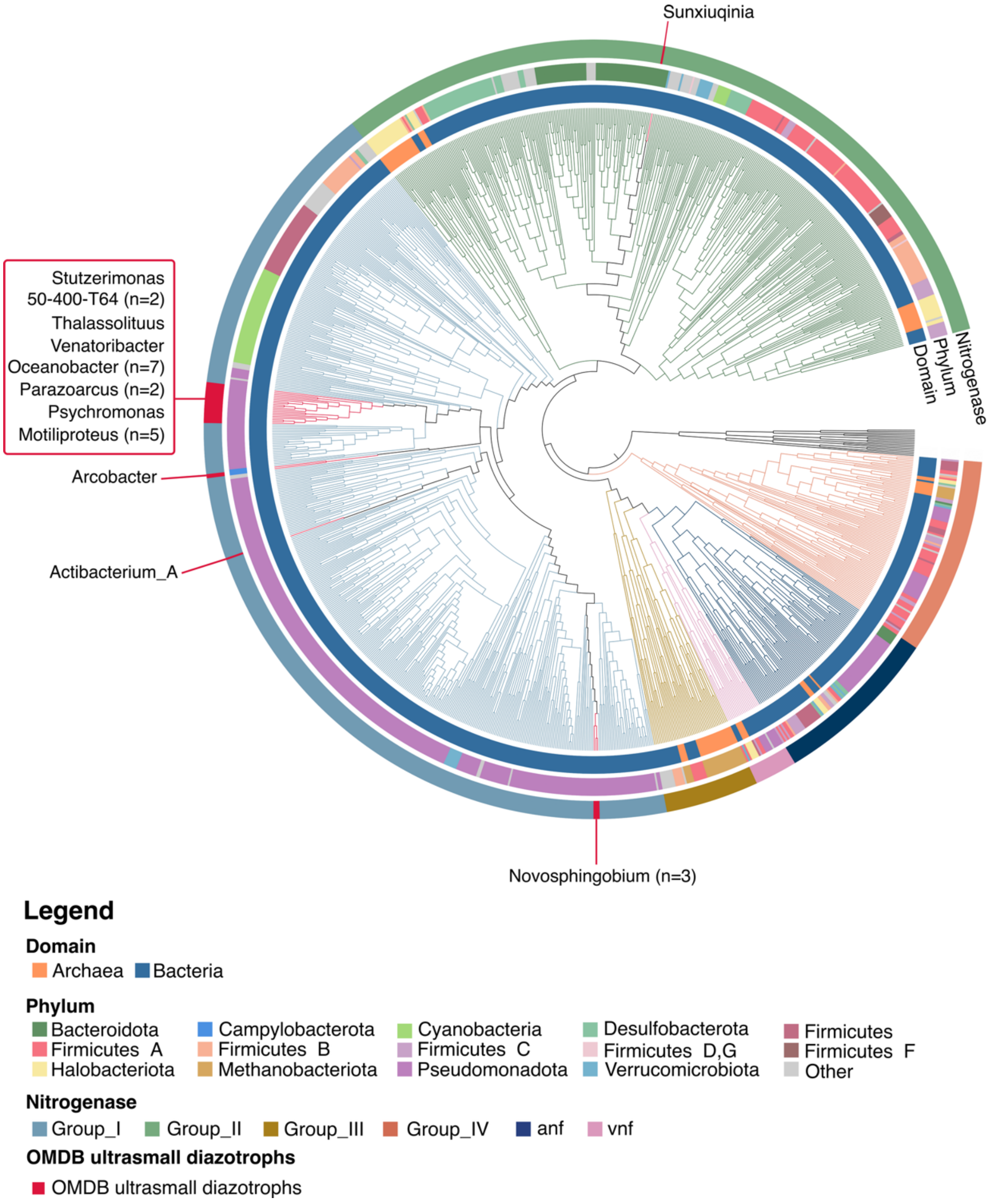


**Figure S7. Phylogeny of concatenated *nifHDK* nitrogenase proteins from diazotroph MAGs from ultrasmall size fractions with respect to reference sequences.** The phylogeny was constructed using concatenated amino acid sequences of *nifHDK* (or *vnfHDK/anfHDK*) from 1,099 reference sequences compiled by Pi et al. (2024)[5], along with 28 MAGs from this study that possess the complete *nifHDK* operon. The tree was inferred using the LG+R10 substitution model and rooted with concatenated outgroup sequences from Bch/ChlLNB and BchXYZ proteins, following the approach of Pi et al. (2024)[5]. The tree is annotated with three concentric rings: the innermost ring indicates domain (Bacteria or Archaea); the second ring shows phylum-level taxonomy; and the outermost ring displays the nitrogenase functional cluster (Groups I, II, III, etc.). MAGs specific to this study are highlighted in red on the outermost ring.

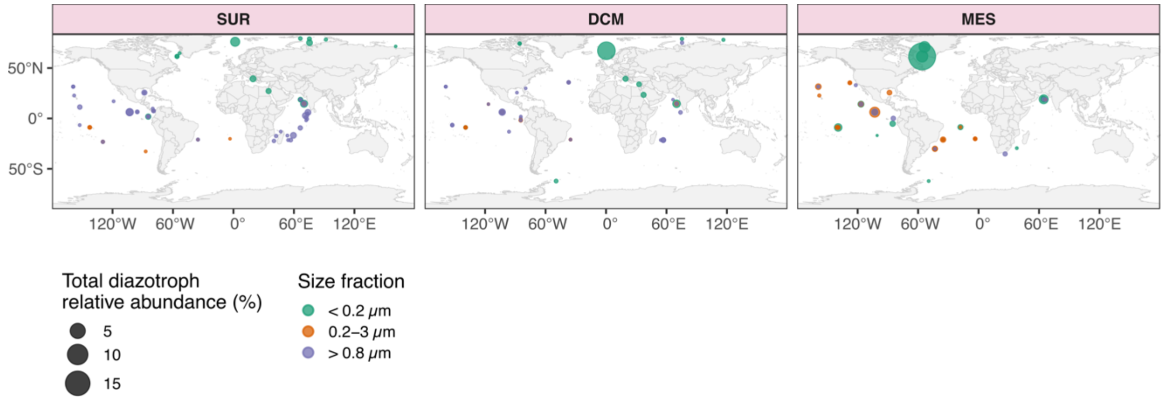


**Figure S8. Distribution of potential diazotrophs by depth.** Depths are indicated respectively as SUR: Surface, DCM: Deep Chlorophyll Maximum, MES: Mesopelagic. Dot size indicated relative abundance of diazotrophs. Color indicated size fraction group.

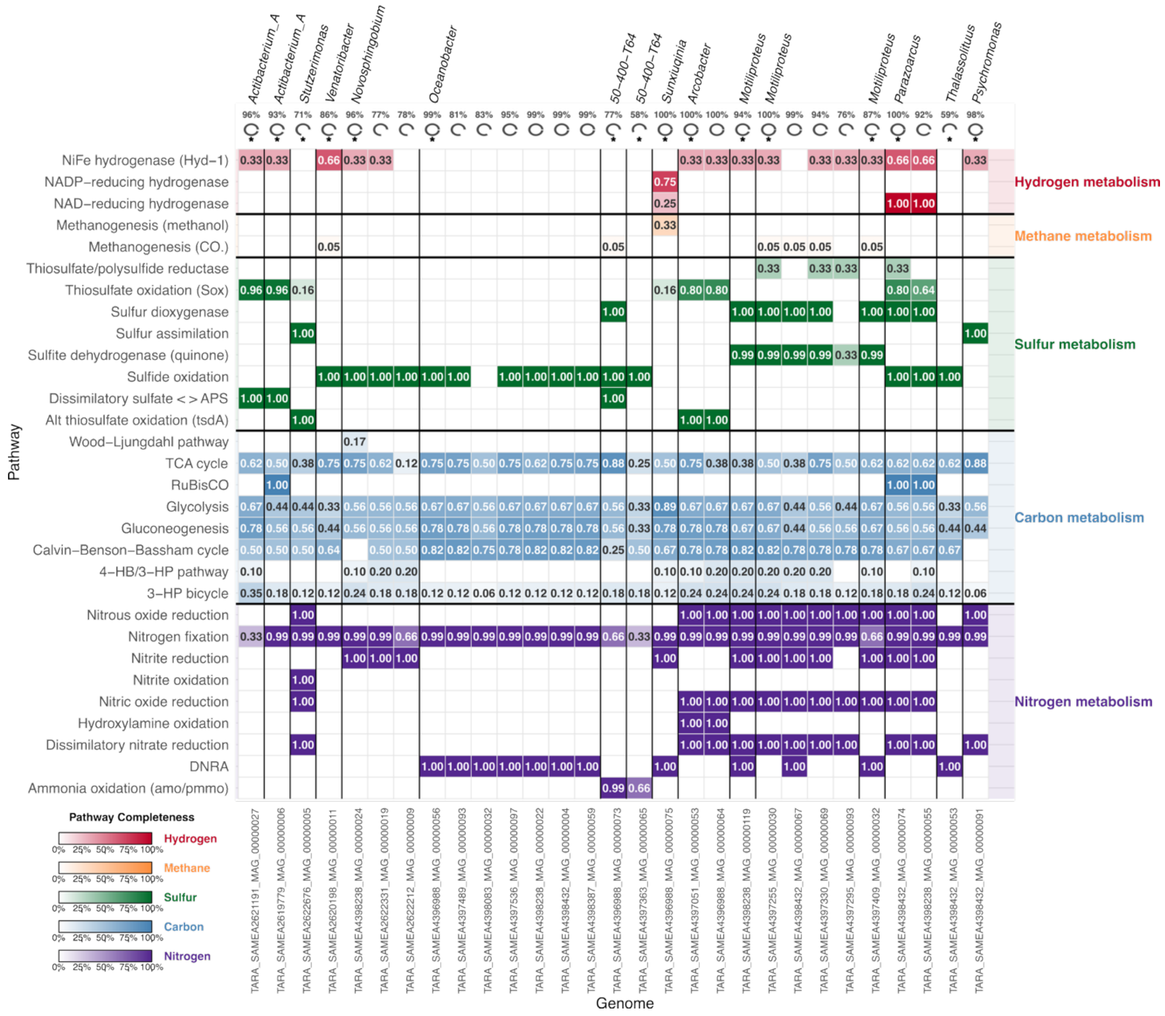

**Figure S9. Metabolic potential of all diazotroph MAGs from ultrasmall size fractions (<0.2 µm).** Key metabolic functions related to hydrogen, methane, sulfur, carbon, and nitrogen cycling of diazotroph MAGs from the ultrasmall size fraction. MAGs are grouped by genus and 99% ANI clusters are separated by black lines. Pathway completeness was determined using KEGG Decoder[6] based on KO annotations and is indicated in the fields. At the top, MAG completeness (%) is shown, with an asterisk (*) marking cluster representatives. On the right, pathways are grouped by metabolic category: hydrogen (red), methane (orange), sulfur (green), carbon (blue), and nitrogen (purple). Color intensity and number in the square indicates the completeness of each KEGG pathway.

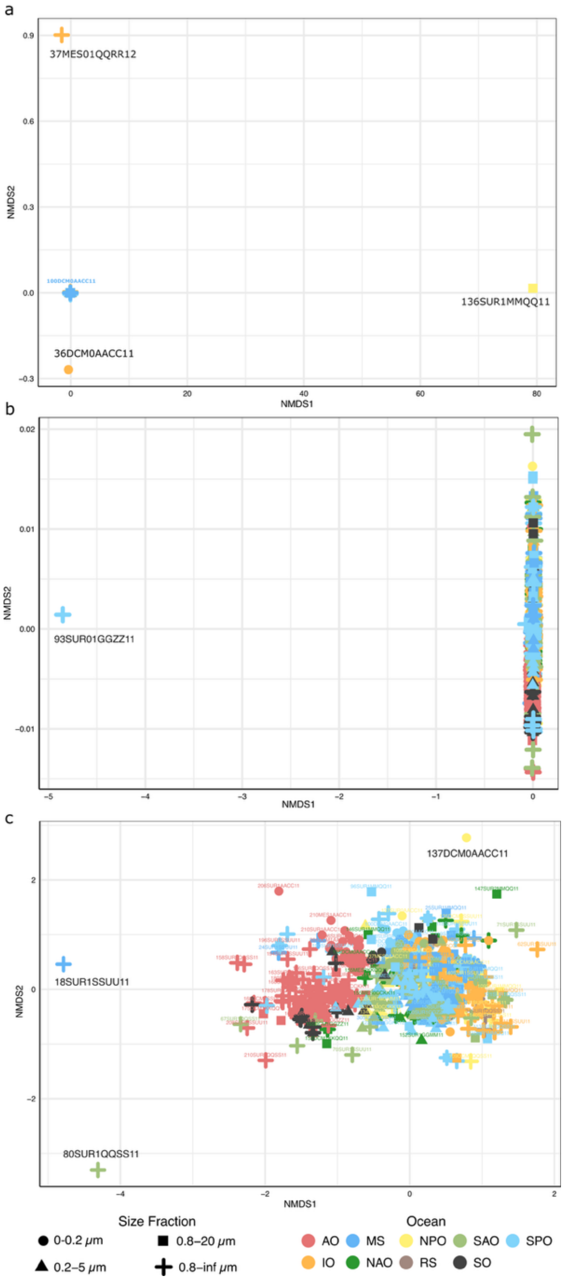


**Figure S10. Non-metric multidimensional scaling (NMDS) based on Bray-Curtis dissimilarity of MAG abundance profiles.** Each point represents a sample, shaped by size fraction and colored by ocean basin. Outliers are written in black **a.** Initial NMDS showing the first three outliers: 37MES01QQRR12, 36DCM0AACC11, and 136SUR1MMQQ11. **b.** NMDS after removal of panel A outliers, revealing sample 93SUR01GGZZ11 as single outlier. **c.** NMDS after removal of panel B outlier, revealing the final three outliers: 18SUR1SSUU11, 80SUR1QQSS11, and 137DCM0AACC11.

**Table S1. Source studies contributing MAGs to the ultrasmall size fraction dataset.** Description of the nine studies from which MAGs were retrieved from the ultrasmall size fraction, including the Ocean Microbiomics Database (OMDB) project accession number, the number of MAGs contributed by each study, the sampling region, and the corresponding reference. These studies span diverse marine environments and depth layers.

| Project | Number of MAGs | Region | Study |
|---|---|---|---|
| ASPL16-1 | 3 | Baltic Sea | Asplund-Samuelsson et al. 2016 [7] |
| DUAR15-1 | 305 | Mediterranean Sea, South Atlantic Ocean, Southern Ocean | Duarte 2015 [8] |
| HEVR20-1 | 56 | Indian Ocean, Red Sea | Hevroni et al. 2020 [9] |
| JAHN19-1 | 2 | Mediterranean Sea | Jahn et al. 2009 [10] |
| LINN22-1 | 69 | North Pacific Ocean | Linney et al. 2022 [11] |
| LUOE20-1 | 169 | North Pacific Ocean | Luo et al. 2020 [12] |
| NILS19-1 | 241 | Baltic Sea | Nilsson et al. 2019 [1]3 |
| RUSA22-1 | 61 | Arctic Ocean | Rusanova et al. 2022 [14] |
| TOPC | 3152 | Mediterranean Sea, Red Sea, Indian Ocean, South Atlantic Ocean, North Atlantic Ocean, North Pacific Ocean, South Pacific Ocean, Arctic Ocean | Sunagawa et al. 2020 [15] |

**Table S2. PERMANOVA analysis of MAG community structure.** Community composition was analyzed using PERMANOVA on Bray-Curtis dissimilarities with 10,000 permutations. MAG presence was estimated when ≥60% of the MAG was covered in reads. NMDS stress = 0.131.

a. **Overall PERMANOVA Results**

| Factor | df | $R^2$ | F | p-value |
|---|---|---|---|---|
| Ocean basin | 8 | 0.087 | 13.62 | < 0.0001 |
| Size fraction | 3 | 0.051 | 21.45 | < 0.0001 |
| Depth | 2 | 0.026 | 16.34 | < 0.0001 |
| Residual | 1045 | 0.832 | - | - |
| **Total** | 1058 | 1.000 | - | - |

b. **Pairwise PERMANOVA: Ocean Basin -** Top 10 Strongest Effects (by $R^2$)

| Ocean 1 | Ocean 2 | $R^2$ | F | P-value (FDR-adj) |
|---|---|---|---|---|
| Southern Ocean | Red Sea | 0.134 | 8.49 | <0.0001 |
| Arctic Ocean | Mediterranean Sea | 0.105 | 33.28 | <0.0001 |
| Arctic Ocean | Indian Ocean | 0.098 | 38.30 | <0.0001 |
| Southern Ocean | Mediterranean Sea | 0.094 | 12.79 | <0.0001 |
| Arctic Ocean | South Pacific Ocean | 0.092 | 40.41 | <0.0001 |
| Arctic Ocean | North Pacific Ocean | 0.090 | 28.79 | <0.0001 |
| Arctic Ocean | North Atlantic Ocean | 0.084 | 26.18 | <0.0001 |
| Arctic Ocean | South Atlantic Ocean | 0.079 | 26.49 | <0.0001 |
| Southern Ocean | North Atlantic Ocean | 0.072 | 9.72 | <0.0001 |
| Southern Ocean | North Pacific Ocean | 0.072 | 10.24 | <0.0001 |

c. **Pairwise PERMANOVA: Size Fraction**

| Size Fraction 1 | Size Fraction 2 | $R^2$ | F | P-value (FDR-adj) |
|---|---|---|---|---|
| 0–0.2 µm (ultrasmall) | 0.2–5 µm | 0.055 | 31.42 | <0.0001 |
| 0–0.2 µm (ultrasmall) | 0.8–20 µm | 0.054 | 16.59 | <0.0001 |
| 0–0.2 µm (ultrasmall) | 0.8–inf µm | 0.039 | 21.02 | <0.0001 |
| 0.2–5 µm | 0.8–20 µm | 0.036 | 20.13 | <0.0001 |
| 0.2–5 µm | 0.8–inf µm | 0.031 | 24.60 | <0.0001 |
| 0.8–20 µm | 0.8–inf µm | 0.09 | 4.81 | <0.0001 |

*Note : The ultrasmall fraction showed the strongest differentiation from all other size fractions*

**Table S3. Shannon diversity** between A. ultrasmall and larger size fractions. B. Ocean basin.

**a. Comparison of species-level Shannon diversity (H′) between ultrasmall and larger size fractions**

| Ocean Basin | Region | Ultrasmall (0–0.2 µm) | Other fractions (>0.2 µm) | ΔH′ |
|---|---|---|---|---|
| Southern Ocean (SO) | Polar | 2.84 ± 0.13 | 0.75 ± 0.40 | 2.10 |
| Arctic Ocean (AO) | Polar | 3.06 ± 0.70 | 1.39 ± 0.48 | 1.67 |
| North Atlantic Ocean (NAO) | Non-Polar | 3.30 ± 0.10 | 2.10 ± 0.68 | 1.20 |
| South Atlantic Ocean (SAO) | Non-Polar | 2.89 ± 0.33 | 2.07 ± 0.76 | 0.82 |
| Mediterranean Sea (MS) | Non-Polar | 2.38 ± 0.43 | 1.74 ± 0.61 | 0.64 |
| Indian Ocean (IO) | Non-Polar | 2.58 ± 0.63 | 1.99 ± 0.87 | 0.59 |
| Red Sea (RS) | Non-Polar | 2.36 ± 0.80 | 1.91 ± 0.93 | 0.45 |
| South Pacific Ocean (SPO) | Non-Polar | 2.42 ± 0.53 | 2.06 ± 0.70 | 0.36 |
| North Pacific Ocean (NPO) | Non-Polar | 2.21 ± 1.27 | 2.14 ± 0.84 | 0.08 |

**b. Mean ± Standard Deviation Shannon Diversity by Ocean Basin**

| Ocean Basin | Species-level H′ | Phylum-level H′ | Region |
|---|---|---|---|
| Arctic Ocean (AO) | 1.78 ± 0.89 | 0.51 ± 0.22 | Polar |
| Southern Ocean (SO) | 1.17 ± 0.93 | 0.52 ± 0.21 | Polar |
| North Atlantic Ocean (NAO) | 2.12 ± 0.88 | 0.85 ± 0.32 | Non-Polar |
| South Atlantic Ocean (SAO) | 2.20 ± 0.88 | 0.70 ± 0.27 | Non-Polar |
| North Pacific Ocean (NPO) | 2.14 ± 0.95 | 0.68 ± 0.29 | Non-Polar |

| | | | |
|---|---|---|---|
| South Pacific Ocean (SPO) | 2.10 ± 0.88 | 0.66 ± 0.27 | Non-Polar |
| Indian Ocean (IO) | 2.13 ± 0.95 | 0.73 ± 0.30 | Non-Polar |
| Mediterranean Sea (MS) | 1.80 ± 0.79 | 0.48 ± 0.19 | Non-Polar |
| Red Sea (RS) | 2.05 ± 0.97 | 0.76 ± 0.30 | Non-Polar |

**Table S4. Summary of the 17 strain level cluster representatives (99% ANI) of potential diazotrophs.** Bold MAGs with a * indicate cluster representatives at ANI 95%.

| MAG | Size | Completeness | *nifH* primer compability | nif gene presence | GTDB-based taxonomy | ANI match | Cluster size (99% ANI) |
|---|---|---|---|---|---|---|---|
| **TARA_SAMEA4396988_MAG_00000056*** | 4085924 | 98.54 | X nifH4 | ✓ | g__Oceanobacter;s__Oceanobacter sp913061185 | Arc-Gamma-03 (99.37) | 7 MAGs |
| **TARA_SAMEA4398238_MAG_00000119*** | 4218063 | 94.37 | X nifH4 | ✓ | g__Motiliproteus;s__ | Arc-Gamma-04 (99.77) | 1 MAG |
| TARA_SAMEA4397255_MAG_00000030 | 4103094 | 100 | X nifH4 | ✓ | g__Motiliproteus;s__ | Arc-Gamma-04 (98.29) | 4 MAGs |
| TARA_SAMEA4397409_MAG_00000032 | 3494936 | 87.32 | X nifH4 | ✓ | g__Motiliproteus;s__ | Arc-Gamma-04 (98.12) | 1 MAG |
| **TARA_SAMEA4398238_MAG_00000024*** | 4431845 | 96.34 | - | ✓ | g__Novosphingobium;s__Novosphingobium indicum | TARA_AON_82_MAG_00070 (98.49) | 3 MAGs |
| **TARA_SAMEA2621191_MAG_00000027*** | 3420314 | 95.82 | No nifH | X *nifH* | g__Actibacterium_A;s__Actibacterium_A naphthalenivorans | - | 1 MAG |
| TARA_SAMEA2619779_MAG_00000006 | 3386494 | 93.12 | - | ✓ | g__Actibacterium_A;s__Actibacterium_A naphthalenivorans | - | 1 MAG |
| **TARA_SAMEA4396988_MAG_00000073*** | 2506789 | 77.44 | X nifH4 | X *nifE* | g__50-400-T64;s__50-400-T64 sp913058345 | - | 1 MAG |
| TARA_SAMEA4397363_MAG_00000065 | 1932708 | 57.75 | X nifH 1,2,4 | X *nifN* | g__50-400-T64;s__50-400-T64 sp913058345 | - | 1 MAG |
| **TARA_SAMEA4397051_MAG_00000053*** | 3019296 | 100 | - | ✓ | g__Arcobacter;s__ | Arc-Campylo (99.99) | 2 MAGs |
| **TARA_SAMEA4398432_MAG_00000074*** | 4691050 | 100 | X nifH4 | ✓ | g__Parazoarcus;s__Parazoarcus communis_A | - | 2 MAGs |
| **TARA_SAMEA4398432_MAG_00000053*** | 3038015 | 58.62 | X nifH4 | X *nifN* | g__Thalassolituus;s__ | - | 1 MAG |
| **TARA_SAMEA2620198_MAG_00000011*** | 3753208 | 86.47 | X nifH4 | ✓ | g__Venatoribacter;s__Venatoribacter sp002706025 | TARA_ION_MAG_00014 (99.97) | 1 MAG |
| **TARA_SAMEA4398432_MAG_00000091*** | 4630116 | 98.05 | X nifH4 | ✓ | g__Psychromonas;s__ | Arc-Gamma-02 (99.62) | 1 MAG |
| **TARA_SAMEA2622676_MAG_00000005*** | 3384469 | 70.98 | X nifH4 | ✓ | g__Stutzerimonas;s__Stutzerimonas stutzeri_G | - | 1 MAG |
| **TARA_SAMEA4396988_MAG_00000075*** | 4619685 | 100 | X nifH4 | X *nifN* | g__Sunxiuqinia;s__ | Arc-Bactero (99.83) | 1 MAG |
| **RUSA22-1_SAMN23283061_MAG_00000043*** | 2011355 | 89.89 | - | X *nifD* | g__JAIOSF01;s__ | - | 1 MAG |

### Supplementary Datasets

**Dataset S1 (separate file).** Supplementary Data Tables. This Excel file contains seven tables:

Supplementary Data 1, ultrasmall MAGs metadata from the Ocean Microbiomics Database (OMDB);

Supplementary Data 2, environmental metadata for all samples analyzed in this study;

Supplementary Data 3, dereplicated MAG clusters at 95% ANI;

Supplementary Data 4, results from NMDS ordination and PERMANOVA statistical analyses; Supplementary Data 5, distribution of MAGs across different size fractions;

Supplementary Data 6, RuBisCO gene (K01601) detection and annotation data;

Supplementary Data 7, curated *nif* gene annotations and classifications based on HMM profiles;

Supplementary Data 8, KEGG orthology assignments for nif genes.

**Dataset S2 (separate file).** Amino acid sequences of *nif* genes identified in ultrasmall MAGs.

**Dataset S3 (separate file).** Nucleotide sequences of *nif* genes identified in ultrasmall MAGs.

**Dataset S4 (separate file).** KEGG Decoder output showing functional pathway completeness across ultrasmall MAGs.

**Dataset S5 (separate file).** Amino acid sequences of *RuBisCO* genes (K01601) identified in ultrasmall MAGs.
